# Feedforward regulatory logic underlies robustness of the specification-to-differentiation transition and fidelity of terminal cell fate during *C. elegans* endoderm development

**DOI:** 10.1101/2021.08.24.457588

**Authors:** Chee Kiang Ewe, Erica M. Sommermann, Josh Kenchel, Sagen E. Flowers, Morris F. Maduro, Joel H. Rothman

## Abstract

Development is driven by gene regulatory networks (GRNs) that progressively dictate specification and differentiation of cell fates. The architecture of GRNs directly determines the specificity and accuracy of developmental outcomes. We report here that the core regulatory circuitry for endoderm development in *C. elegans* is comprised of a recursive series of interlocked feedforward modules linking a cascade of six sequentially expressed GATA-type transcription factors. This structure results in a reiterated sequential redundancy, in which removal of a single factor or alternate factors in the cascade results in no, or a mild, effect on endoderm development and gut differentiation, while elimination of any two factors that are sequentially deployed in the cascade invariably results in a strong phenotype. The strength of the observed phenotypes is successfully predicted by a computational model based on the timing and levels of transcriptional states. The feedforward regulatory logic in the GRN appears to ensure timely onset of terminal differentiation genes and allows rapid and robust lockdown of cell fate during early embryogenesis. We further found that specification-to-differentiation transition is linked through a common regulator, the END-1 GATA factor that straddles the two processes. Finally, we revealed roles for key GATA factors in establishing spatial regulatory state domains by acting as transcriptional repressors that appear to define the boundaries of the digestive tract. Our findings support a comprehensive model of the core gene network that describes how robust endoderm development is achieved during *C. elegans* embryogenesis.

**Graphic abstract:** 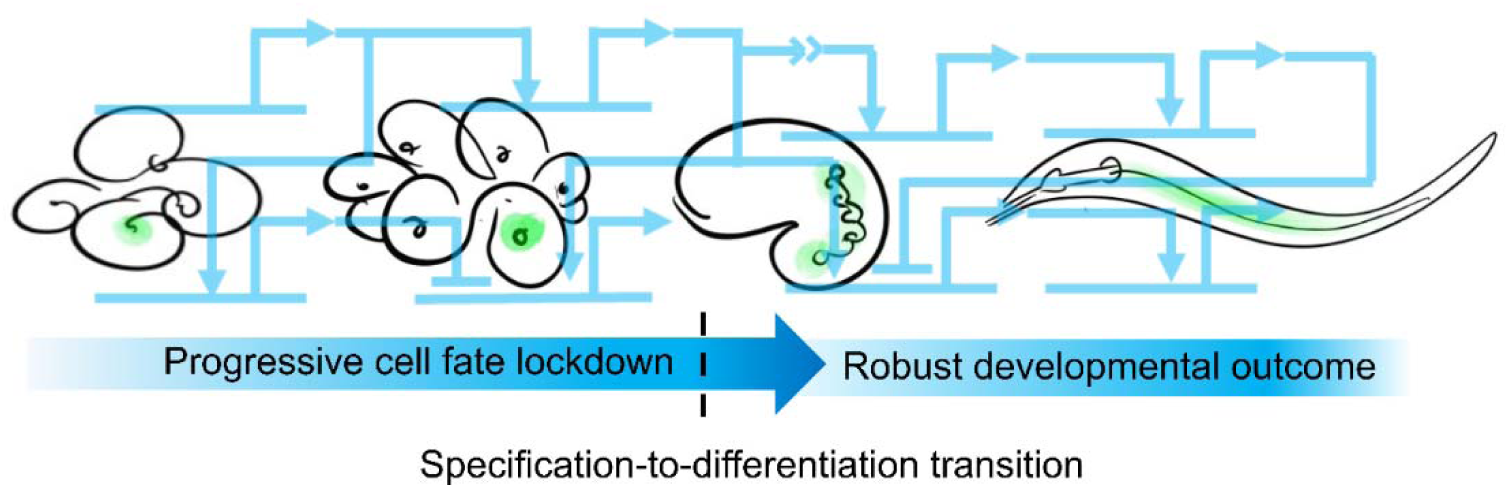

## Introduction

Development is driven by progressive activation of transcriptional programs, or gene regulatory networks (GRNs) that direct cell specification and subsequent differentiation (Boveri, 1899; Davidson and Levine, 2008). The sequential restriction of cell identity and developmental potential through dynamic changes in transcriptional states was anticipated by Waddington’s epigenetic landscape, a graphic metaphor describing canalization and robustness during development (Waddington, 1957).

The generation of diverse animal forms largely relies on a common genetic toolkit. The GATA transcription factors play a conserved role in the development of diverse cell types, including those of the endoderm, the first of the three germ layers to have evolved during the late Precambrian era (Hashimshony et al., 2015; Rodaway and Patient, 2001). In the diploblastic phyla Poriferans and Cnidarians, GATA factors have been found to be specifically expressed in the endoderm, suggesting that these transcriptional regulators may have driven the invention of the endoderm germ layer and gastrulation at the dawn of metazoan evolution (Martindale et al., 2004; Nakanishi et al., 2014). This association of GATA factors with endoderm development persists throughout metazoan phylogeny, generally through successive deployment of multiple GATA factors. This sequential use of GATA factors is particularly striking in *C. elegans*, in which a cascade of six GATA factor-like transcription factors regulate specification and differentiation of the endoderm (Maduro, 2017; Maduro and Rothman, 2002; McGhee, 2007). In *Drosophila*, the GATA factor Serpent specifies endodermal fate and activates the expression of a second GATA factor, dGATAe, which is essential for the terminal differentiation of the intestine. These factors can act across wide phylogenetic spans, as first demonstrated with the *C. elegans* END-1 GATA factor, which ectopically activates endoderm development when expressed in the prospective ectoderm of *Xenopus* (Shoichet et al., 2000). Similarly, overexpression of *serpent* or *dGATAe* causes ectopic endoderm differentiation in non-endodermal lineages in *Drosophila*, as well as in *Xenopus*, further supporting the functional conservation of the GATA factors (Murakami et al., 2005; Okumura et al., 2005). In sea urchin, Blimp1/Krox1 activates *otx1*, the product of which activates *gatae* expression (Davidson et al., 2002). Gatae in turn provides a positive input to *otx1*, in addition to activating the transcriptional program for endoderm development, thereby forming a stable circuit in the GRN (Davidson et al., 2002; Yuh et al., 2004). Accordingly, knocking down *gatae* severely blocks endoderm development and gastrulation (Davidson et al., 2002).

The endoderm in *C. elegans*, which arises from a single blastomere born at the 8-cell stage, the E cell (Boveri, 1899; Sulston et al., 1983), provides a highly tractable system for investigating the mechanisms of cell specification, differentiation, and organogenesis. This progenitor cell gives rise to a clone 20 cells comprising the intestine, which are arranged in nine rings (int1-9) spanning the length of the animal (Supplementary Figure 1A). Endoderm development is driven by pairs of duplicated genes encoding GATA-like transcription factors: the divergent MED-1/2 factors and the canonical END-1/3 and ELT-2/7 factors. The maternally provided SKN-1/Nrf transcription factor activates MED-1 and -2, which activate the specification of mesendodermal fate in the EMS blastomere (Bowerman et al., 1992). In the anterior daughter of EMS, the MS cell, the Wnt effector POP-1/Tcf represses the expression of *end-1/3*, and MED-1 and -2 activate *tbx-35*, whose product specifies mesodermal fate. In the E cell, the posterior daughter of EMS, a triply redundant signaling system (Wnt, Src, and MAPK) leads to the phosphorylation of POP-1 by nemo-like kinase LIT-1 (Bei et al., 2002; Maduro et al., 2002; Meneghini et al., 1999; Shin et al., 1999; Thorpe et al., 1997). Together with MED-1/2, Wnt-signaled POP-1 activates genes encoding the transiently expressed endoderm specification factors END-1 and -3, which in turn activate the expression of *elt-7* and *-2*, orthologues of vertebrate *GATA4*/*5/6*. Expression of the ELT factors is sustained throughout the remainder of the animal’s life via a positive autoregulatory loop that “locks down” the differentiated state of the intestine (Supplemental Figure 1B) (Maduro, 2017; Maduro and Rothman, 2002; McGhee, 2007). While *elt-7* loss-of-function mutants do not show a discernible phenotype, animals lacking ELT-2 arrest at early larvae stage (L1) owing to severely obstructed gut (Sommermann et al., 2010). Nonetheless, *elt-2(-)* mutant animals contain a well-defined intestinal lumen and the intestinal cells appear to be fully differentiated (Fukushige et al., 1998; Sommermann et al., 2010). In the absence of both ELT-2 and -7, however, the intestinal lumen is completely abolished and differentiation appears to proceed in only a subset of the endoderm-derived cells in a sporadic manner (Sommermann et al., 2010). This suggests that ELT-2 and -7 function synergistically in mediating morphological differentiation of the intestine, and additional input(s) mediate a bistable switch in the endodermal differentiation program in the absence of ELT-2 and -7 (Dineen et al., 2018; Sommermann et al., 2010).

In this study, we sought to decipher the functional requirements and interactions between the GATA transcription factors and how they allow rapid and faithful deployment of the endoderm GRN. We found that the endoderm GRN consists of a series of interlocking feedforward loops, creating “sequential redundancy” in the cascade and culminating in the rapid lockdown of cell fate. We further report that END-1 acts at a transition point, participating in both specification and differentiation. Finally, we demonstrated the important roles of GATA factors in safeguarding intestinal cell fate and defining the boundaries of the digestive tract. Overall, our findings reveal the nature of the extensive genetic redundancy in the regulatory circuitry and how this GRN architecture dictates robust cell specification and differentiation during embryonic development.

## Methods and Materials

### *C. elegans* cultivation and genetics

Worm strains were cultured using standard procedure (Brenner, 1974) and all experiments were performed at room temperature (20-23°C). All genetic manipulations were performed according to standard techniques (Fay, 2013). *him-5(-)* or *him-8(-)* was introduced into some strains to generate males and facilitate crosses. See Supplementary Table 1 for a complete list of strains used in this study.

### Immunofluorescence analysis

Antibody staining with methanol-acetone fixation was performed as previously described (Sommermann et al., 2010). Antibodies MH27 (AB_531819), MH33, and 455-2A4 (AB_2618114) were used to detect AJM-1, IFB-2, and ELT-2, respectively. Alexa Fluor® 488 goat anti-mouse secondary antibody was used at 1:1000 dilution.

### RNAi

RNAi feeding clones were obtained from the Ahringer (Kamath et al., 2003) or the Vidal (Rual et al., 2004) library. The bacterial strain was inoculated overnight at 37°C in LB containing 50 μg/ml ampicillin. The bacterial culture was then diluted 1:10 and incubated for an additional 4 h. Next, 1 mM of IPTG was added to the bacterial culture and 100 μL was seeded onto 35 mm NGM agar plates containing 1 mM IPTG and 25 μg/mL carbenicillin. For simultaneous knockdown of *elt-2* and *elt-7*, the two bacterial strains, each expressing dsRNA for one gene, were concentrated and resuspended in 1 mL of LB in 1:1 ratio before seeding the NGM plates. Seeded plates were allowed to dry for 48 h before use. Next, 10-20 L4 animals were placed on the RNAi plates. 24 h later, the animals were transferred to fresh RNAi plates to lay eggs. The progeny was then collected for analyses.

### Imaging and fluorescence quantification

The animals were immobilized using 10 mM Levamisole and mounted on 4% agarose pads. Images were acquired, typically at 60X, using Nikon Eclipse Ti-E inverted microscope fitted with ORCA-Flash2.8 camera. For expression studies, maximum intensity Z-projection was generated on the Nikon NIS-Elements AR v4.13.05. Images were then analyzed using ImageJ or Imaris v9.7.2.

### RT-qPCR

RNA was extracted from synchronized L1 animals using Monarch^®^ Total RNA Miniprep Kit (#TS010S). cDNA synthesis was performed using SuperScript™ III First-Strand Synthesis SuperMix (Thermo Fisher Scientific, #18080400). Quantitative PCR was performed using BioRad CFX96 Real-Time System. Each 15 µL reaction contained cDNA, primers, and PowerUp^TM^ SYBR^TM^ Green Master Mix (Thermo Fisher Scientific, #A25743). The data were analyzed using the standard 2^-ΔΔ^ method (Livak and Schmittgen, 2001).

The primers sequences are:

*act-1* - 5’-TCCATTGTCGGAAGACCACG-3’ and 5’-GGTGACGATACCGTGCTCAA-3’
*act-5* - 5’-GTCACTCACACCGTTCCAATC-3’ and 5’-GTGAGGATCTTCATCATGTAGTCG-3’

### Modeling endoderm gene regulatory circuits

The topology of the gene circuits with temporal information was written as a system of differential equations, with expression of each factor dependent on the concentration of its activators (See Supplementary Materials and Supplementary File 1). The gene cascade is initiated by SKN-1, which was modelled as a square wave in the EMS blastomere. Similarly, the positive inputs of (phosphorylated) POP-1 into *end-3* and *end-1* were modelled as a square wave in the E blastomere (23 mins after the four-cell stage). Model runs were calculated as time-discretized Euler approximations (Hahn, 1991) with time steps of 0.01 s. An iterative least-squares algorithm following a modified Gauss-Newton method (Ruhe, 1979; Yip et al., 2010) was used to fit the model parameters to published transcriptomics data (Baugh et al., 2003; Tintori et al., 2016). The performance of the final model was then evaluated by comparing the predicted phenotypes of the single mutants with published results (Boeck et al., 2011; Dineen et al., 2018; Maduro et al., 2005a; Maduro et al., 2015). Finally, predictions of *elt-2* activation (a readout for the commitment to E fate) in the mutant combinations were generated by holding the concentration of knockout genes at zero and otherwise running the model as described. The source code is available on https://github.com/RothmanLabCode/endoderm_GRN_model.

### Statistics and figure preparation

Statistics were performed using R software v3.4.1 (https://www.r-project.org/). The specific statistical tests were reported in the figure legends. Plots were generated using R package ggplot2 or Microsoft Excel. Figures were assembled in Inkscape v0.92.4 (https://inkscape.org/).

## Results

### Sequential redundancy suggests feedforward regulatory circuitry in the endoderm GRN

Owing to the extensive genetic redundancy, single mutants of most of the genes throughout the *C. elegans* endoderm GRN either show no overt phenotype (single *med* mutants, *end-1(-)*, *elt-7(-)*) or an extremely weakly penetrant phenotype (*end-3(-)*). Moreover, while the *elt-2(-)* mutant undergoes larval arrest immediately after hatching with a dysfunctional gut, the gut appears fully differentiated in these mutants. A strong gut differentiation defect requires the removal of both *elt-2* and *elt-7* (Figure 1A) (Fukushige et al., 1998; Sommermann et al., 2010). Similarly, strong phenotypes are observed only when both *meds* or both *ends* are removed in pairs. In this study, we sought to examine the basis for this extensive redundancy in the pathway and to illuminate how it might contribute to faithful specification and differentiation in endoderm GRN.

**Figure 1:**
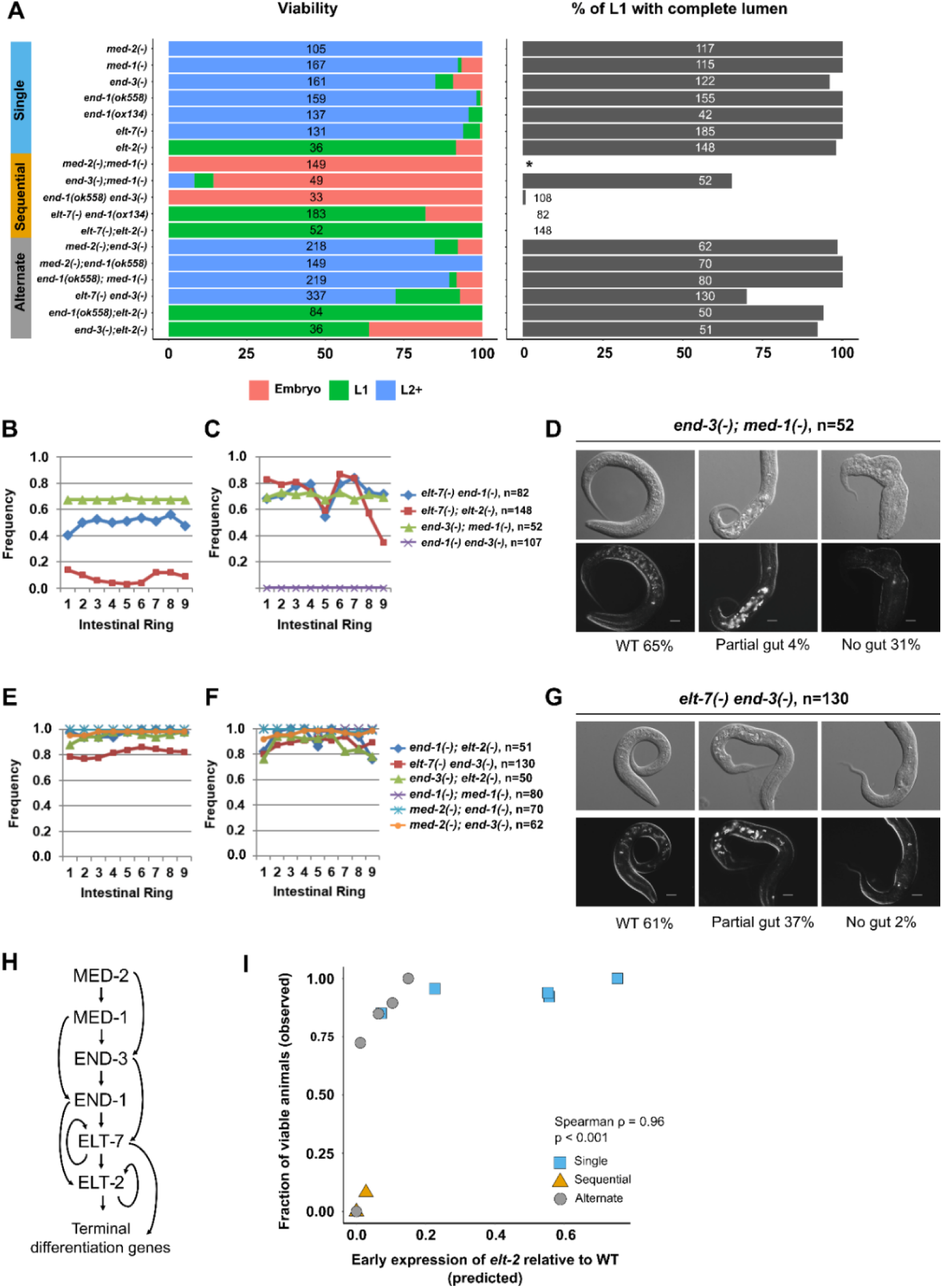
Evidence for recursive feedforward loops in the endoderm GRN. (A) Comparison of viability and gut lumen morphology in mutant combinations lacking one or two GATA factors in the endoderm regulatory cascade. Mutants missing sequential members of the endoderm cascade exhibit more penetrant developmental defects than mutants with alternate steps disrupted. We note the two loss-of-function alleles of *end-1*, *ox134* (14822 bp deletion; also removes the adjacent *ric-7* gene) and *ok558* (879 bp deletion), do not result in significant loss in viability (Fisher’s exact test; p = 0.31) or other discernible phenotypes. We used *ox134* in subsequent analyses unless stated otherwise (see Supplementary Table 1). The total number of animals scored for each genotype is indicated. No L1 animals were scored for *med-2(-); med-1(-)* (indicated by *) as all double mutants arrest as embryos owing to misspecification of MS (and E in majority but not all of the embryos). The frequencies of (B, E) visible lumen and (C, F) gut granules are dramatically reduced in double mutants missing (B, C) sequential GATA pairs compared to those lacking (E, F) alternate members of the endoderm cascade. (D, G) *end-3(-); med-1(-)* shows a more severe defect in intestinal differentiation than *elt-7(-) end-3(-)*. 31% of *end-3(-); med-1(-)* and 2% of *elt-7(-) end-3(-)* show no overt signs of endoderm differentiation. (G) Many *elt-7(-) end-3(-)* larvae contain a partial gut as observed by DIC microscopy and birefringent gut granules, presumably due to sub-threshold levels of END-1-dependent activation of *elt-2*. Scale bar = 10 μm. (H) The sequential redundancy model of the endoderm GRN. (I) Computed *elt-2* expression levels predict the phenotype severity of the mutant combinations. All *med-2(-); med-1(-)* mutants show severe embryonic lethality despite *elt-2* activation owing to a fully penetrant MS *1(-)* from this analysis.

Temporally resolved transcriptomic analyses revealed that the GATA transcription factors are sequentially activated in the mesendoderm regulatory pathway (Baugh et al., 2003; Tintori et al., 2016). This observation suggests that, rather than pairs of factors (MEDs, ENDs, ELTs) acting together at specific tiers in the cascade (Supplementary Figure 1B), each factor is redundant with its immediate upstream or downstream factor in a continuously sequential cascade (“sequential redundancy”). For example, as is often seen in forward-driven biological switches, each may act through a feedforward motif in which a given factor activates both its immediate target gene and the target of that gene (Mangan and Alon, 2003).

To test the model that endoderm differentiation is directed by a series of sequentially redundant feedforward regulatory motifs, we constructed a series of double mutants and compared the penetrance of lethality and the extent of gut differentiation. We found that unlike the mild or undetectable phenotypes observed with single mutants, removing pairs of genes that are expressed at sequential steps in the cascade invariably results in severely diminished viability (Figure 1A) and defects in endoderm specification/differentiation, as revealed by disrupted gut lumen morphology (Figure 1A, B). This effect is associated with sporadic expression and in some cases the complete absence of gut-specific rhabditin granules (Clokey and Jacobson, 1986) (Figure 1C). We found that the expression of the gut-specific IFB (Intermediate <u>F</u>ilament, <u>B</u>)-2 protein is strongly diminished in the double mutants (Supplementary Figure 2A). Moreover, reduced AJM (<u>A</u>pical <u>J</u>unction <u>M</u>olecule)-1 expression suggests gut epithelialization defects in double mutants of sequential members in the endoderm GRN (Supplementary Figure 2B).

In *med-2(-); med-1(-)* double mutants, all embryos die as embryos as MS blastomere fails to be specified and its fate is transformed to that of the C-like mesectodermal progenitor in the early embryo (Figure 1A), as is also the case in embryos lacking the initiating, maternally provided SKN-1 transcription factor (Bowerman et al., 1992). The E cell similarly adopt a C cell fate in some, but not all, of the *med-2(-); med-1(-)* arrested embryos (Maduro et al., 2007). As previously reported (Maduro et al., 2005a), many *end-1(-) end-3(-)* mutants also die as embryos owing to transformation of the E cell into a C-like cell (Figure 1A). Thus, END-1 and -3 act as redundant selector genes that promote specification of endoderm over other cell fates.

Consistent with the sequential redundancy model, removal of two genes that function at alternate steps in the endoderm cascade does not result in the strong synergistic effect that we observed with removal of pairs of sequentially expressed genes (Figure 1A, E, F; Supplementary Figure 3). We observed normal expression of IFB-2 and AJM-1 along the length of the gut region in all such double mutants (Supplementary Figure 3). Further, while the *end-3(-); med-1(-)* double mutant, in which sequential genes in the cascade are removed, shows severe embryonic lethality, most (89.5%; n = 219) of the *end-1(-); med-1(-)* double mutants, in which alternate genes in the cascade are removed, develop into fertile adults (Figure 1A). This observation is also consistent with the reported divergent roles of END-1 and -3 in endoderm specification (Boeck et al., 2011). Moreover, the *elt-7(-) end-3(-)* double mutants are largely viable, in contrast to the *end-3(-); med-1(-)* sequential double mutants, which show a strongly penetrant embryonic lethal phenotype (Figure 1A). Of the hatched L1 larvae, 31% of *end-3(-); med-1(-)* sequential double mutant animals exhibit no overt signs of gut differentiation, while only 2% of *elt-7(-) end-3(-)* alternate double mutants completely lack gut (Fisher’s exact test; p < 0.001) (Figure 1D, G). However, many *elt-7(-) end-3(-)* mutants contain a partially differentiated gut (Figure 1G). We reasoned that the developmental defects observed in the most affected animals are the result of suboptimal expression of *end-1* in the absence of *end-3* (Maduro et al., 2007).

While neither single mutant shows a discernible phenotype, animals lacking both END-1 and ELT-7 are 100% inviable, and the arrested L1 larvae show a striking gut differentiation defect (Figure 1A, Supplementary Figure 2). The mutant worms contain patches of apparently differentiated gut as evidenced by expression of gut granules and immunoreactive IFB-2, similar to *elt-7(-); elt-2(-)* double mutants, which exhibit an all-or-none block to differentiation event along the length of the animals (Figure 1B, C; Supplementary Figure 2) (Sommermann et al., 2010). However, the differentiation defect observed in *elt-7(-) end-1(-)* double mutants appears to be somewhat milder from that in *elt-7(-); elt-2(-)*: unlike the latter, we observed defined, albeit sporadic, lumen and brush border, and the undifferentiated patches are more frequently interspersed with differentiated patches (Figure 1B, C; Supplementary Figure 2A) (more below).

Analysis of most alternate and sequential double mutant combinations show similar effects (Fig. 1A): sequential double mutants are invariably much more severely affected than alternate double mutants. Together, our data support a sequentially redundant cascade, comprising a recursive series of feedforward regulatory steps (Figure 1H). It is conceivable that such a feedforward system creates a strongly forward-driven, rapidly deployed switch that ensures timely and robust cell fate commitment and robust lockdown of endoderm cell fate during embryogenesis.

### Computational model predicts phenotypes of sequential and alternative double mutants

We took a complementary approach to testing the sequential feedforward model through a computational strategy. We constructed a mathematical model based on the network topology, in which the interactions between the GATA factors, as well as the additional POP-1-dependent activation of *end-3* and *end-1*, were written as a series of ordinary differential equations and the model parameters (Supplementary File 1) were determined by fitting to published transcriptomics data (Baugh et al., 2003; Tintori et al., 2016) using a custom algorithm that follows an iterative least-squares method. We then carried out *in silico* perturbations of the feedforward circuits (see Methods and Materials and Supplementary Materials) to provide predictions of these effects on the relative timing and levels of *elt-2* activation and therefore the final output of the endoderm GRN. Our computed results revealed that *elt-2* expression is predicted to occur, but with delayed onset in all single mutants and in mutants lacking alternate pairs of GATA factors (Figure 1I; Supplementary File 1). However, as observed with the experimental outcomes, *elt-2* expression is predicted to be completely abrogated in most double mutant combinations in which sequential members of the endoderm cascade are removed (Figure 1I), consistent with their pronounced developmental defects (Figure 1A). The predicted *elt-2* expression level strikingly correlates (Spearman’s Rank Correlation ρ = 0.96; p < 0.001) with the experimentally observed penetrance of the phenotypes of the single and multiple mutants (Figure 1I): single and alternate double mutant combinations, which show no, or weak phenotypes, are predicted to express high levels of *elt-2* early, while sequential double mutants, which show strong defects in gut development and inviability, are predicted not to express *elt-2* at substantial levels at its normal time of onset. These findings, based on modeling with gene expression data, bolster the proposed recursive feedforward structure of the GATA factor cascade.

### Variation in temporal expression explains distinct functions for MED-1 and -2

It has been shown previously that *end-3* expression is activated slightly earlier than *end-1* (Supplementary Figure 4) (Maduro et al., 2007; Tintori et al., 2016), as is also consistent with our genetic analyses (Figure 1). Additionally, the END paralogues have diverged considerably with variable DNA-binding domains that perform overlapping but distinct functions (Boeck et al., 2011). Unlike the ENDs, MED-1 and -2 are 98% identical (Maduro et al., 2001). Nonetheless, our genetic evidence demonstrates distinguishable contributions of the two nearly identical paralogues, with MED-2 functions preceding MED-1 (Figure 1; Supplementary Figure 2, 3). By examining published lineage-resolved single-cell RNA-seq data (Tintori et al., 2016), we observed that *med-2* transcripts are undetectable by the 8-cell stage, while *med-1* expression persists briefly in the endoderm precursors (Supplementary Figure 4A). The differential temporal expression of the MEDs was recapitulated using protein-fusion reporters: unlike MED-1, which is expressed in 16E embryos, MED-2 protein expression is largely diminished by the 8E embryonic stage (Supplementary Figure 4B). Together, our data suggest *med-1* and *-2* genes are differentially regulated and MED-2 functions upstream of MED-1, further supporting the feedforward structure at the beginning of the endoderm regulatory cascade.

### Synergistic requirements and cross-regulatory interactions of END-1, ELT-7, and ELT-2

As described above, while both *elt-7(-)* and *elt-2(-)* single mutants contain a fully differentiated gut with a contiguous lumen from the pharynx to the rectum surrounded by cells of normal differentiated morphology, *elt-7(-); elt-2(-)* double mutants invariably lack both a defined gut lumen as well as some gut cells, and show a sporadic, all-or-none, block to gut differentiation along the length of the animals (Figure 1A-C; Figure 2A; Supplementary Figure 2, 5) (Sommermann et al., 2010). Although differentiation is highly defective in the absence of ELT-2 and -7, patches of well-differentiated gut are nonetheless evident. Moreover, many terminal differentiation genes remain activated in the absence of ELT-2 and -7 (Dineen et al., 2018). For example, eliminating the functions of ELT-2 or -7 has little effect on the expression of *act-5*, a gene encoding an actin isoform required for microvilli formation (Dineen et al., 2018; MacQueen et al., 2005). These observations suggest that at least one additional factor, in addition to the ELTs, may activate gut differentiation. One such candidate is END-1, which acts in specification of E cell identity immediately upstream of the *elt* genes. Indeed, although *end-1(-)* and *elt-7(-)* mutants are both phenotypically silent, we found that the *elt-7(-) end-1(-)* double mutant shows extensive gut differentiation defects, with sporadic expression of rhabditin granules (Figure 1A-C; Figure 2B) and *ifb-2* (Figure 2C, D; Supplementary Figure 2), as well as reduced number of differentiated gut cells (Figure 2E; Supplementary Figure 6), reminiscent of *elt-7(-); elt-2(-)* double mutant animals.

**Figure 2:**
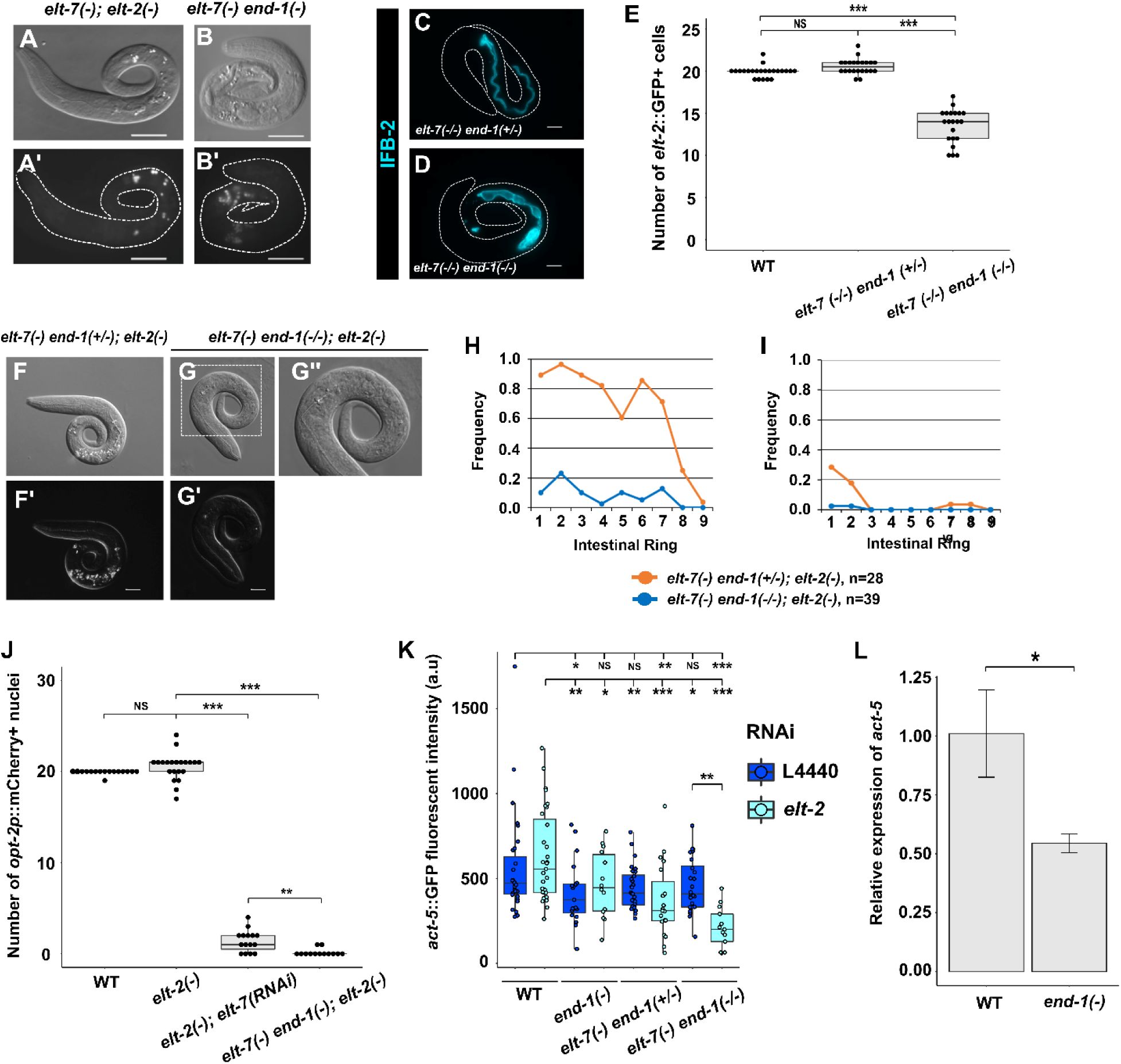
Synergistic actions of END-1, ELT-7 and ELT-2 mediate morphological differentiation of endoderm. (A, B) *elt-7(-) end-1(-)* contains a grossly defective gut with sporadic patches of rhabditin granules interspersed with apparently undifferentiated regions, similar to that observed in *elt-7(-); elt-2(-)*. Scale bars = 20 μm. (C, D) Balanced *elt-7(-/-) end-1(-/+)* larva shows uniform expression of IFB-2 (*kcIs6* transgene) along the length of the animal, while *elt-7(-/-) end-1(-/-)* shows sporadic expression of the transgene. Scale bars = 10 μm. (E) Wildtype and *elt-7(-/-) end-1(-/+)* animals contain an average of 20 intestinal cells, marked by *elt-2*::GFP reporter. The number of differentiated gut cells is markedly reduced in *elt-7(-/-) end-1(-/-)* larvae (mean = 13.5 cells). (F, G) Representative micrographs of *elt-7(-) end-1(+/-); elt-2(-)* and *elt-7(-) end-1(-/-); elt-2(-)* triple mutants. (F, F’) *elt-7(-) end-1(+/-); elt-2(-)* lacks a visible lumen and contains sporadic birefringent granules. (G, G’) *elt-7(-) end-1(-/-); elt-2(-)* exhibits absolutely no signs of endoderm differentiation. (G”) Magnified view of the animal present in (G). Scale bars = 10 μm. (H, I) The average frequencies of (H) gut granule and (I) lumen are dramatically reduced in *elt-7(-) end-1(-/-); elt-2(-)* compared to *elt-7(-) end-1(+/-); elt-2(-)* animals. (J) Wildtype and *elt-2(-)* show similar number of differentiated intestinal cells, although the variance in *elt-2(-)* is significantly increased (F-test p < 0.001). The number of *opt-2p::mCherry* (*irSi24*) expressing cells is strongly reduced in *elt-2(-); elt-7(RNAi)*. No gut cells were detected in vast majority of *elt-7(-) end-1(-); elt-2(-)* triple mutants. (K) The expression of *act-5::GFP* translational reporter *(jyIs13*) in various mutant combinations. END-1, ELT-7, and ELT-2 appear to act together to regulate *act-5* expression in the intestine. (L) *act-5* is downregulated in *end-1(-)* compared to wildtype as detected by RT-qPCR. *act-1* was used as the internal reference. Three replicates were performed for each genotype. Error bars represent standard deviation. * p ≤ 0.05 by two-tail t-test. For panels E and J, NS p > 0.05, ** p ≤ 0.01, *** p ≤ 0.001 by non-parametric Kruskal-Wallis test and pairwise Wilcoxon tests with Benjamini & Hochberg correction. For panel K, NS p > 0.05, * p ≤ 0.05, ** p ≤ 0.01, *** p ≤ 0.001 by parametric one-way ANOVA followed by pairwise t-tests with Benjamini & Hochberg correction.

We further tested whether END-1 could account for the residual gut-promoting activity by removing it from animals also lacking ELT-2/7. Strikingly, simultaneously eliminating END-1, ELT-7, and ELT-2 results in an apparent elimination of intestinal differentiation (Figure 2F-I). We found that 19.6% of *elt-7(-) end-1(-); elt-2(-)* mutants undergo embryonic arrest, and the remainder die as L1 larvae (n = 143). The triple mutant animals exhibit no morphological signs of gut differentiation, showing no gut granules (Figure 2F, G, H), no detectable intestinal brush border or lumen (Figure 2F, G, I), and no differentiated gut nuclei expressing a gut-specific peptidase transporter, OPT-2 (Figure 2J; Supplementary Figure 7), suggesting a near total block to gut differentiation. We found that knocking out *end-1* strongly reduces the expression level of *act-5* (Figure 2K, L), and *act-5* expression is further downregulated in *elt-7(-) end-1(-)* and *elt-7(-) end-1(-); elt-2(RNAi)* animals (Figure 2K; Supplementary Figure 8), suggesting that END-1, ELT-7 and ELT-2 act collaboratively and redundantly to mediate *act-5* expression, and that END-1 may compensate for the loss of ELT-2 and -7 inputs.

A challenge to the notion of a role for END-1 as the gut-promoting factor acting in the absence of the ELTs is that its expression in wildtype embryos is transient, such that its product is largely undetectable by the 16E embryonic stage (Li et al., 2019; Zhu et al., 1997); yet gut differentiation seen in the differentiated patches in the *elt-7(-); elt-2(-)* double mutant appears strong and robust late in development. These findings led us to hypothesize that ELT-2 and/or ELT-7 may normally repress *end-1* transcription through negative feedback and that in the absence of the ELTs, *end-1* may be upregulated and drive differentiation. Indeed, we found that the intensity of an *end-1* endogenous protein fusion reporter is modestly elevated in 8E embryos when *elt-2* is knocked down by RNAi (Figure 3A). Moreover, the expression level of *end-1* is upregulated in both 4E and 8E embryos when ELT-7 and -2 are simultaneously depleted (Figure 3B). While these findings provide evidence for the hypothesized feedback inhibition, the effect is rather weak, as we did not observe obvious longer-term perdurance of *end-1* expression (see Discussion).

**Figure 3:**
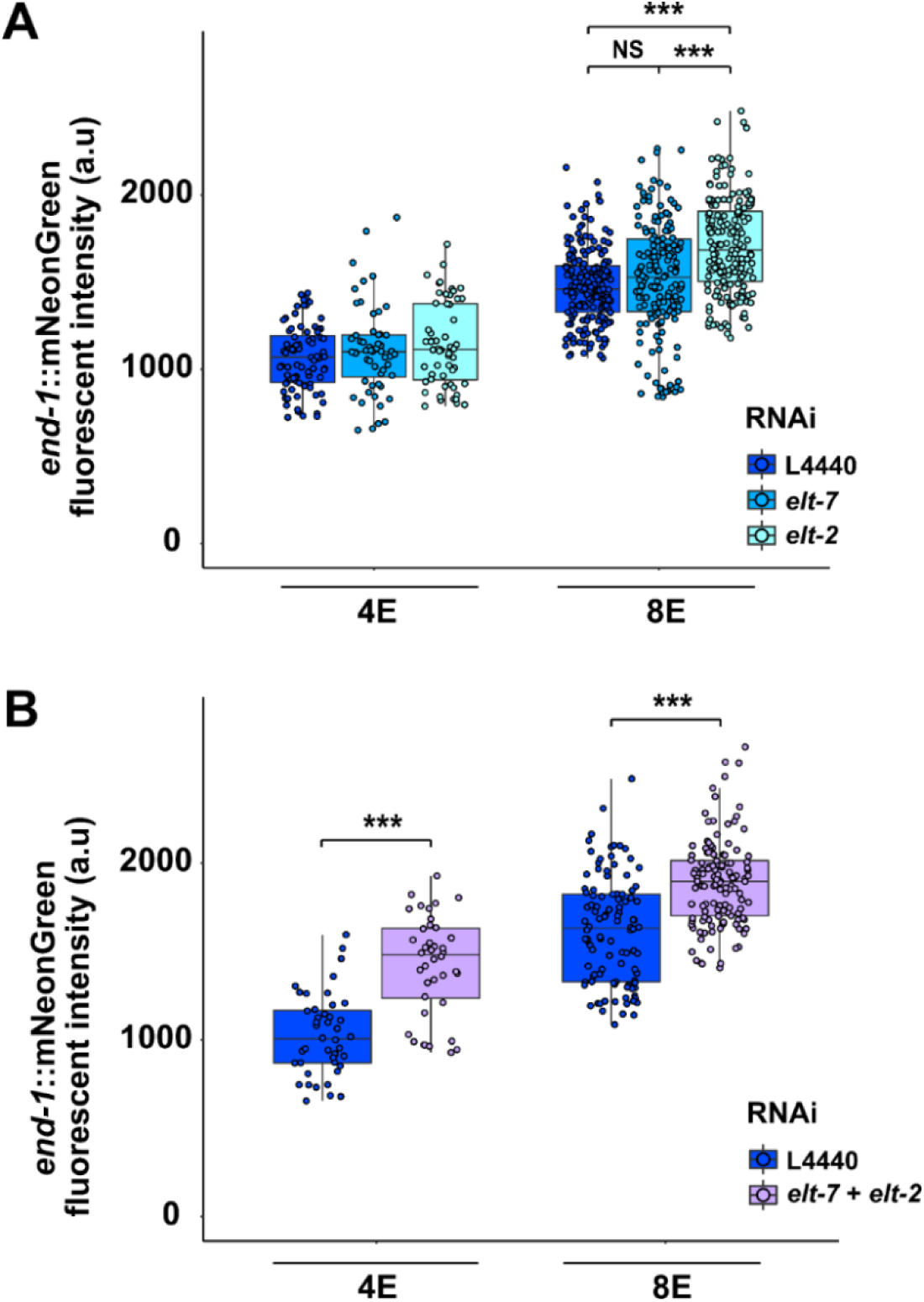
ELT-2 antagonizes *end-1* expression. (A) Expression of endogenously tagged *end-1* increases in 8E embryos upon *elt-2* RNAi treatment. NS p > 0.05, *** p ≤ 0.001 by Kruskal-Wallis test and pairwise Wilcoxon tests with Benjamini & Hochberg correction. (B) Knocking down both *elt-7* and *elt-2* further elevates END-1 expression in both 4E and 8E embryos. *** p ≤ 0.001 by Wilcoxon tests.

Taken together, our results suggest END-1 is poised in the cascade at the interface between specification and differentiation, placing it at the crux of this key transition. END-1, acting with END-3, regulates *specification* of the E lineage, whereas END-1, acting with ELT-7 and ELT-2, controls *differentiation* of the intestine.

### ELT-2 and ELT-7 collaborate to safeguard intestinal cell fate

When improperly specified, E has been shown to undergo wholesale lineage conversion to either a C-like mesectodermal fate (when SKN-1, the MEDs, or END-1/3 are removed), thereby inappropriately generating epidermis, or an MS-like mesodermal fate (for example, in the absence of Wnt signaling), thereby inappropriately giving rise to pharyngeal tissue (details described in Figure 4A). Given that END-1 straddles the transition from specification of the endoderm progenitor and differentiation of the gut, we sought to investigate whether specification and differentiation involve distinct regulatory events by examining non-endodermal (epidermal and pharyngeal) gene activity when differentiation is impaired after this transition is thought to occur. In *end-1(-) end-3(-)* double mutants, misspecification of the E cell and its conversion to a C-like mesectodermal progenitor, results in severe morphological defects as a result of the supernumerary epidermal cells and consequent deformation of the epidermis (Figure 4B; Supplementary Figure 9) (Maduro et al., 2005a). In contrast, the *elt-7(-) end-1(-); elt-2(-)* triple mutant animals that, like the *end-1(-) end-3(-)* double mutants, do not make a discernible gut, are not substantially defective for overall body morphogenesis (Supplementary Figure 9), suggesting that E is initially specified. Consistent with this observation, unlike *end-1(-) end-3(-)* double mutant animals, *elt-7(-) end-1(-); elt-2(-)* triple mutant larvae contain a wildtype number of epidermal cells, implying that E C misspecification does not occur (Figure 4B). These results imply that END-3 alone is sufficient to promote endoderm specification over C-like mesectodermal cell fate. However, we were surprised to observe mis-expression of the pharyngeal muscle-specific myosin gene, *myo-2*, in the gut region of many *elt-7(-); elt-2(-)* and *elt-7(-) end-1(-); elt-2(-)* mutants (Figure 4C, D). Additionally, knocking down *elt-7* in *end-1(-); elt-2(-)* results in ectopic expression of *ceh-22*, which encodes a pharynx-specific NK-2-type homeodomain protein, in the otherwise undifferentiated gut (Supplementary Figure 10). As the Wnt/POP-1-dependent polarization of EMS is unperturbed in the mutants, we reasoned that inappropriate expression of pharyngeal genes is unlikely to be the result of wholesale E→MS transformation, but may reflect later errors in the fidelity of differentiation.

**Figure 4:**
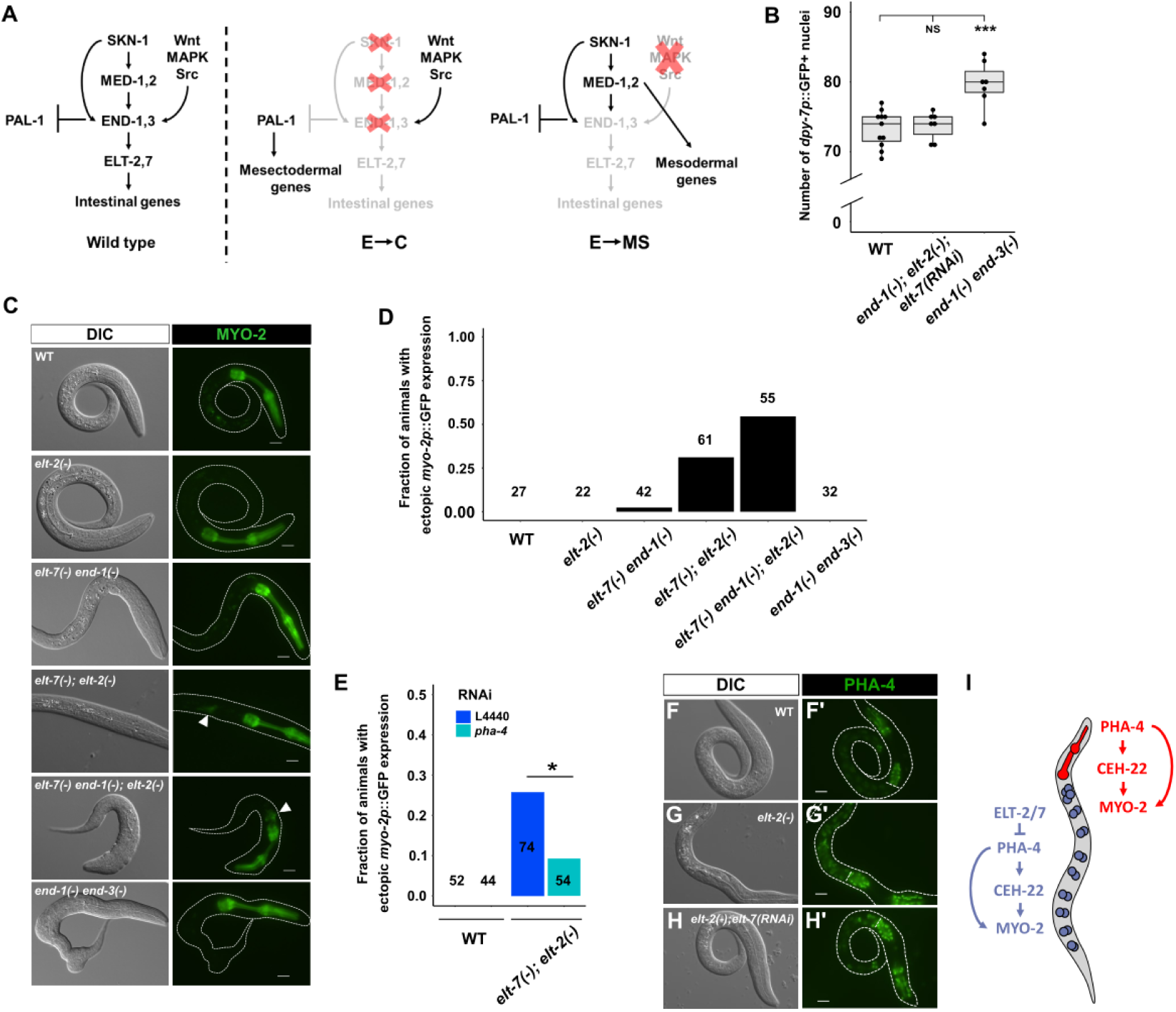
ELT-2 and ELT-7 repress pharyngeal fate in the intestine. (A) E/C or E/MS binary fate choice during early embryogenesis. The Caudal homolog PAL-1 is required for the specification of mesectodermal C blastomere, which gives rise to epidermis and body wall muscles (Hunter and Kenyon, 1996). Maternally provided *pal-1* is specifically translated in the EMS and P_2_ cells. In MS and E, PAL-1 activity is blocked by TBX-35 and END-1/3, respectively (Broitman-Maduro et al., 2006). Thus, depleting TBX-35, END-1/3, or their upstream activators, MED-1/2 and SKN-1, causes excess skin and muscle owing to the misspecification of MS and/or E as C, the somatic descendant of P_2_ (Hunter and Kenyon, 1996). As EMS divides, Wnt, MAPK, and Src signaling from P_2_ polarizes EMS, leading to the phosphorylation and nucleocytoplasmic redistribution of POP-1, and activation of *end-1/3* in E, but not MS. When the polarizing signal from P_2_ is disrupted, POP-1 is unphosphorylated, and MED-1 and -2, instead, activate the development of mesoderm (MS), which gives rise to the posterior pharynx and body wall muscles (Maduro and Rothman, 2002; Maduro et al., 2002; Rocheleau et al., 1999; Shin et al., 1999). (B) Wildtype and *end-1(-); elt-2(-); elt-7(RNAi)* larvae contain ∼73 epidermal cells, while *end-1(-) end-3(-)* larvae contain ∼80 epidermal cells marked by *dpy-7p::GFP* expression. NS p > 0.05, *** p ≤ 0.001 by parametric one-way ANOVA followed by pairwise t-tests with Benjamini & Hochberg correction. (C) Representative DIC and fluorescent micrographs showing different mutation combinations expressing *myo-2p::GFP*. Ectopic expression of *myo-2* is evident in *elt-7(-); elt-2(-)* and *elt-7(-) end-1(-); elt-2(-)* mutants (arrowheads). (D) The frequency of animals showing mis-expression of *myo-2* as shown in (C). Number of animals scored for each genotype is indicated. (E) Knocking down *pha-4* partially rescues ectopic expression of *myo-2* in *elt-7(-); elt-2(-)* animals. * p ≤ 0.05 by Fisher’s exact test. (F-H) The expression of endogenously tagged *pha-4* reporter in (F, F’) wildtype, (G, G’) *elt-2(-)*, and (H, H’) *elt-2(-); elt-7(RNAi)* worms. The white horizontal lines in (F’-H’) mark the posterior end of the pharynx. Exposure time = 195 ms. All scale bars = 10 μm. (I) Model of spatial repression and fate exclusion in the digestive tract. ELT-2 and -7 repress *pha-4* and prevent the expression pharyngeal genes in the midgut.

PHA-4/FoxA is the organ selector gene that specifies pharyngeal identity, regulating *myo-2* and *ceh-22* among thousands of other targets in the pharynx (Mango et al., 1994; Zhong et al., 2010). We found that the ectopic expression of *myo-2* in *elt-7(-); elt-2(-)* is at least partially suppressed in *pha-4(RNAi)* animals (Figure 4E). In wildtype animals, PHA-4 is expressed at high levels in the pharynx and rectum and at low levels in the intestine (Figure 4F, F’) (Horner et al., 1998; Kalb et al., 1998). It was previously found that ELT-2 positively regulates *pha-4* as forced expression of *elt-2* causes widespread activation of *pha-4* (Kalb et al., 1998). Paradoxically, we found that *pha-4* is upregulated in the intestine in *elt-2(-)* animals (Figure 4G, G’), and depleting *elt-7* in *elt-2(-)* animals further enhances this effect (Figure 4H, H’). These findings raise the possibility that ELT-2 serves dual roles as both an activator and a repressor depending on its expression level. Thus, it appears that upregulation of *pha-4* in the midgut in the absence of ELT-2 and -7 activates sporadic ectopic pharyngeal gene activity. Supporting our model, PHA-4 target genes have been shown to be regulated in part by PHA-4 binding affinity and occupancy (Fakhouri et al., 2010; Gaudet et al., 2004). Taken together, our results show that the boundaries of regulator state domains along the digestive tract of *C. elegans* are established, at least partly, by transcriptional repression mediated by ELT-2 and -7 in the intestine (Figure 4I).

### END-1 and ELT-7 establish the boundary between the valve and intestinal tubes

The foregoing results suggest that the core regulators involved in gut differentiation (END-1, ELT-2, and ELT-7) regulate faithful differentiation of cells in the digestive tract. The pharynx and the intestine are linked by the pharyngeal-intestinal valve (vpi), which consists of six cells arranged into three rings (Rasmussen et al., 2013) (Figure 5A). In wildtype worms, *ajm-1*::*GFP* (*jcIs1* transgene) is strongly expressed through the adherens junctions lining the lumen of the pharynx and vpi, and the expression drops off sharply to low levels starting at the anteriormost ring of the intestine, and continuing throughout the entire length of the animal (Köppen et al., 2001; Sommermann et al., 2010). However, while *end-1(-)* and *elt-7(-/-) end-1(+/-)* animals show wildtype *ajm-1* expression pattern, *ajm-1* signal is markedly elevated in the anterior intestinal terminus of *elt-7(-/-) end-1(-/-)* animals (Figure 5B-E). Additionally, we observed ectopic expression of two vpi markers, *cdf (<u>c</u>ation <u>d</u>iffusion facilitator)-1* (Figure 5F-H) and *hum (<u>h</u>eavy chain, <u>u</u>nconventional <u>m</u>yosin)-1* (Figure 5I-K), in the anterior terminus of *end-1(-/-) elt-7(-/-)* larvae.

**Figure 5:**
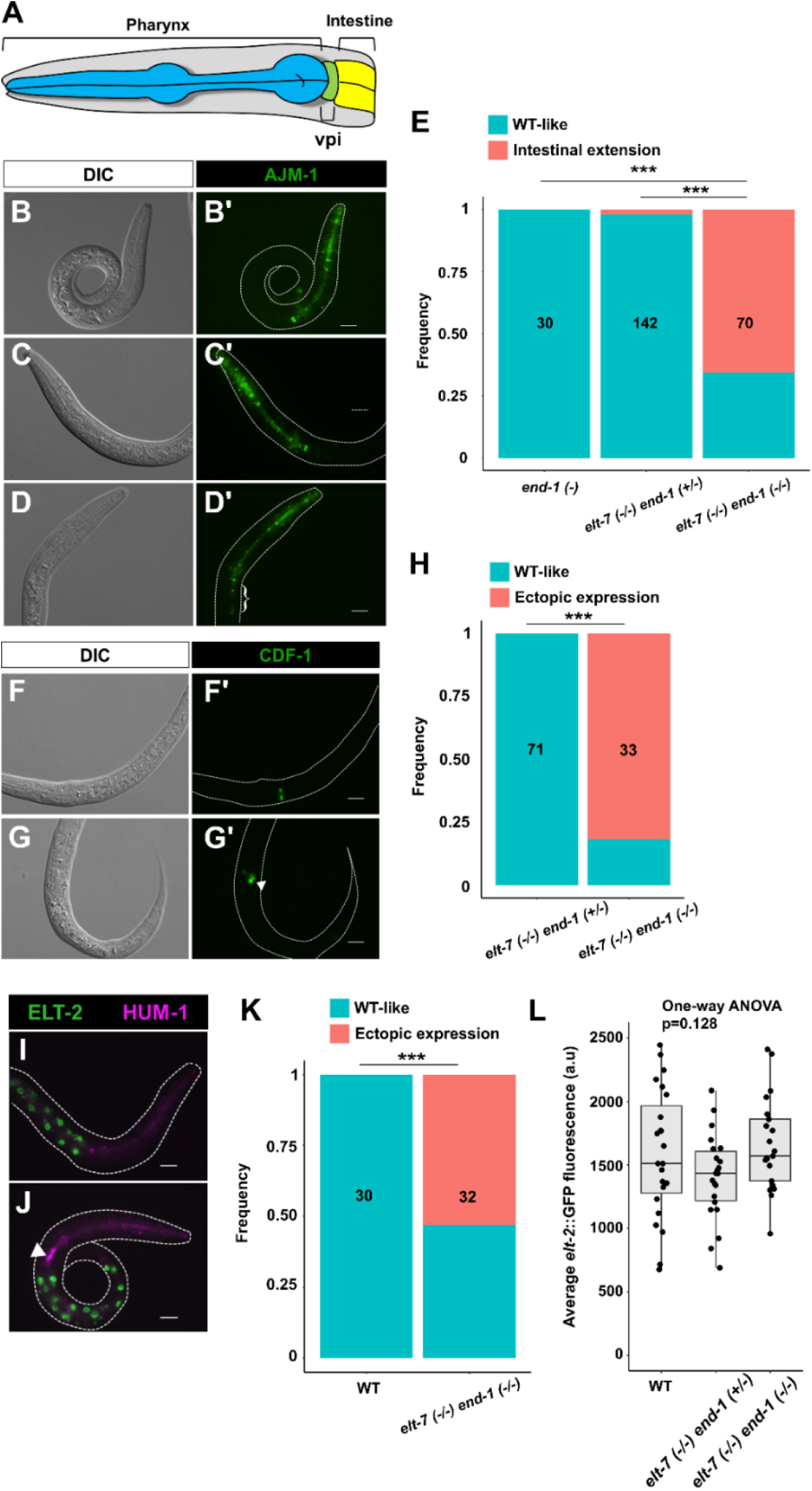
ELT-7 and END-1 function synergistically to repress ectopic expression of valve cell markers in the anterior gut. (A) Drawing highlighting the anatomy of pharynx, vpi, and intestine. The expression of *jcIs1[ajm-1::GFP]* transgene appears wildtype in (B, B’) *end-1(-)* and (C, C’) balanced *elt-7(-/-) end-1(+/-)* mutants. (D, D’) *elt-7(-) end-1(-)* shows caudal expansion of *jcIs1* expression into the anterior gut (bracket). (E) The frequency of animals exhibiting intense *jcIs1* reporter expression in the anterior gut. (F, F’) *cdf-1::GFP* expression is restricted to the vpi in *elt-7(-/-) end-1(+/-)*. (G, G’) Ectopic expression of *cdf-1* reporter is observed in *elt-7(-/-) end-1(-/-)* animals (arrowhead). (H) The frequency of animals with ectopic *cdf-1::GFP* expression in the anterior gut. (I) HUM-1 is highly expressed in the vpi in wildtype animal as revealed by an endogenously tagged reporter. (J) Ectopic expression of *hum-1* is observed in *elt-7(-/-) end-1(-/-)* (arrowhead).

It has been previously shown that worms lacking *elt-2*, like *elt-7(-/-) end-1(-/-)* animals, exhibit striking caudal extension of the valve cell markers (Sommermann et al., 2010). We found that ELT-2 is expressed at wildtype levels in *elt-7(-/-) end-1(-/-)* larvae, owing to its positive autoregulation (Figure 5L). This suggests that the expansion of vpi gene expression in the intestine that we observed in *elt-7(-/-) end-1(-/-)* animals may be independent of ELT-2 function. It is currently unclear whether the expression of vpi reporters in the intestine reflects *bona fide* transformation of gut cells into valve-like cells or aberrant development of vpi and mispositioning of excess valve cells. Regardless, our results demonstrate the important roles of intestinal GATA factors in the development of a properly patterned digestive tract.

The intestinal cells are marked by *elt-2::GFP*. (K) The frequency of animals with ectopic *hum-1::RFP* expression in the anterior gut. (L) The expression of ELT-2 is not altered in *elt-7(-/-) end-1(+/-)* and *elt-7(-/-) end-1(-/-)*, compared to wildtype L1 larvae. All scale bars = 10μm. For panels E, H, K, the number of animals scored is indicated in each graph. *** p ≤ 0.001 by Fisher’s exact test.

## Discussion

The development of *C. elegans* endoderm provides a powerful system to study the regulatory logic underlying cell specification and differentiation. In this study, we reported four major findings: (1) the hierarchical organization and feedforward regulation of GATA factors promote robust and rapid lockdown of endodermal cell fate during *C. elegans* early embryogenesis. (2) END-1 participates in both specification and differentiation and mediates a smooth regulatory state transition. (3) ELT-2 and -7 repress expression of *pha-4* in the midgut to establish regulatory state boundary between the pharynx and the intestine. (4) END-1, ELT-7, and ELT-2 repress the characteristics of pharyngeal valve cell fate at the anterior gut terminus, further defining the spatial domains of the foregut and midgut. Our study therefore provides an important insight into the regulatory circuits that direct specification-to-differentiation transition and subsequent restriction and maintenance of differentiation pattern during development.

### Architecture of the *C. elegans* Endoderm Regulatory Cascade

This study, together with findings reported in other publications, provided a comprehensive view of the core endoderm regulatory cascade, with six GATA factors acting through reiterated sequential feedforward loops (Figure 6). At the top of the cascade, maternally provided SKN-1 binds to and turns on the *meds* and the *ends* (Maduro et al., 2001; Maduro et al., 2005b). Although MED-1 and MED-2 protein sequences are nearly identical, we found distinguishable contributions between the two paralogs, with MED-2 acting upstream of MED-1. Indeed, embryos lacking MED-1 show a weaker loss-of-gut phenotype than those lacking MED-2, when SKN-1 function is debilitated (Maduro et al., 2007). Moreover, *med-2* is expressed slightly earlier than *med-1* (Maduro et al., 2007; Tintori et al., 2016), suggesting they are differentially regulated as we have observed in this study.

**Figure 6:**
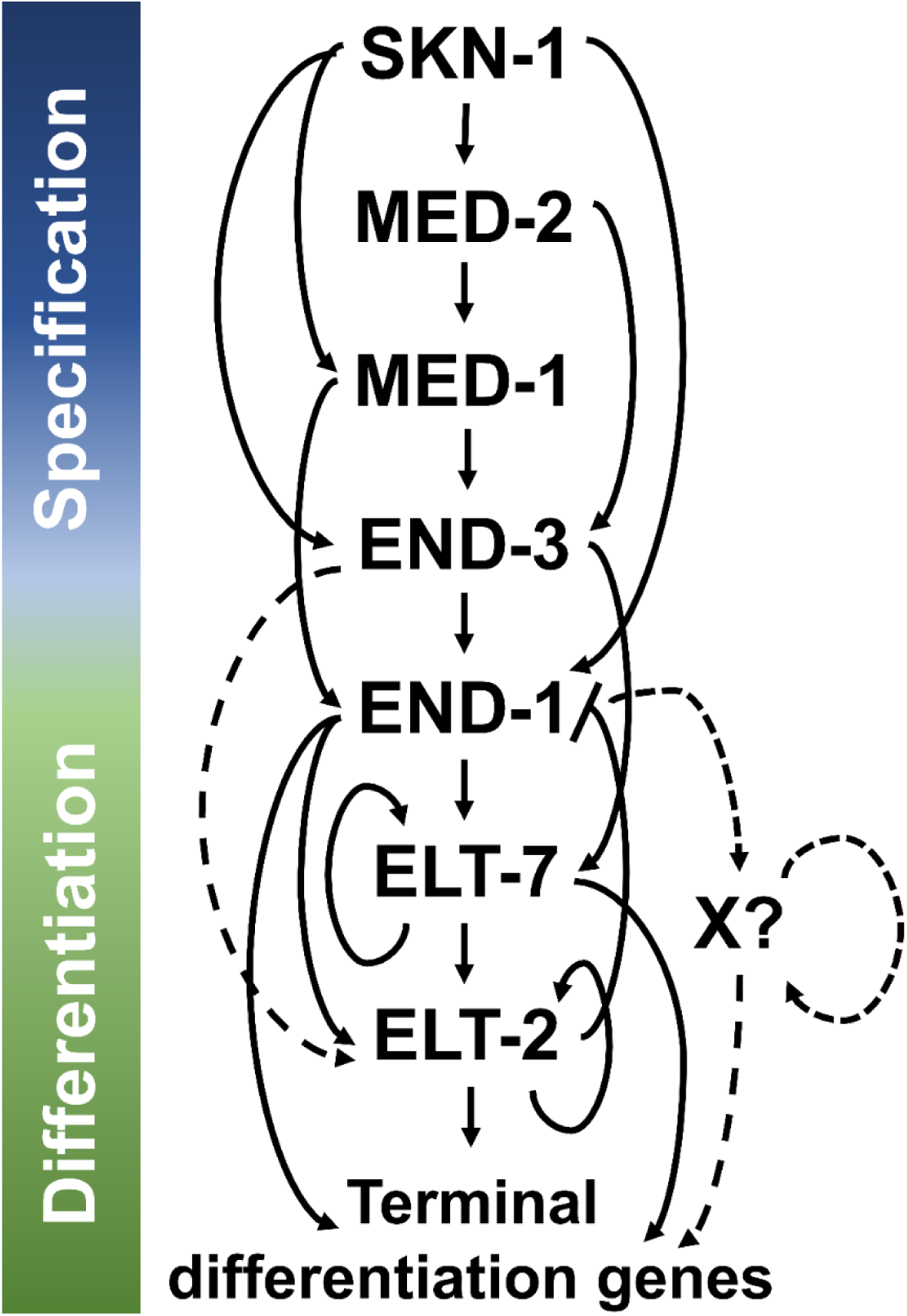
Current model for the *C. elegans* endoderm GRN. Solid lines indicate known interactions identified biochemically or implied genetically, while dashed lines represent proposed interactions. Maternal factor SKN-1 initiates the endoderm cascade which consists of GATA transcription factors arranged in interlocking feedforward circuits. END-1 and -3 are also regulated by non-GATA transcriptional factors, including SPTF-3 (Sullivan-Brown et al., 2016), PAL-1 (Maduro et al., 2005b), PLP-1 (Witze et al., 2009), and POP-1 (Maduro et al., 2005b), which are omitted from this model for simplicity. The observation of residual gut differentiation in *elt-7(-) end-(-)* larvae suggests that END-3 may directly act on ELT-2 and initiate gut differentiation program in some, but not all, of the endodermal progenitors in the double mutants. END-1 may activate an unidentified transcription factor (X) which functions to promote gut differentiation after END-1 decays at ∼16E embryonic stage (see text for details).

In the E blastomere, SKN-1 and the MEDs collaboratively activate END-3. MED-1/2 and END-3 in turn activate END-1, the next step in the cascade. While the two endoderm-specifying factors, END-1 and -3, perform largely overlapping functions, our data suggest that END-3 alone is sufficient to direct endoderm specification and suppress inappropriate mesectodermal development, consistent with the lack of detectable phenotype of in *end-1(-)* single mutants. Interesting, we found that END-1, poised at the interface between specification and differentiation, works synergistically with ELT-2 and -7 to activate the endoderm differentiation program (Figure 6). Supporting our model, in vitro gel-shift assays demonstrate the binding of END-1, ELT-7, and ELT-2 to TGATAA sites, which are highly enriched in the promotors of intestinal genes (Du et al., 2016; McGhee et al., 2009; Wiesenfahrt et al., 2016). Remarkably, END-1 is able to initiate endoderm differentiation in *Xenopus* embryos, demonstrating its role as a potent organ selector (Shoichet et al., 2000). Hence, it appears that specification and differentiation involve a *bona fide* handoff of regulatory events by END-1.

### Regulatory Logic of Developmental GRN

Our results support the conclusion that the *C. elegans* endoderm GRN comprises a recursive series of feedforward steps, culminating with rapid terminal differentiation (Figure 6). Each GATA factor in the entire cascade receives redundant activating inputs acting through an “OR” logic gate. Consequently, any single mutation in the regulatory cascade is largely phenotypically silent, with the exception of *elt-2(-)*; however, even *elt-2(-)* animals contain what appears to be a well-differentiated, albeit dysfunctional, intestine. Coherent feedforward motifs of the type we observe reiteratively in the endoderm GRN are ubiquitous in developmental GRNs. Such a network configuration appears to result in a rapid response to an activating signal and a delayed response when the inputs are removed (termed sign-sensitive delay), thereby prolonging the effect of a transient activator (Mangan and Alon, 2003). Additionally, such feedforward loops are effective at buffering the system against stochastic noise, thereby ensuring developmental robustness (Chepyala et al., 2016; Gui et al., 2016; Maduro, 2015). This design principal appears to be crucial to ensure timely and robust activation of *elt-7* and *-2*, as we and others have shown (Boeck et al., 2011; Dineen et al., 2018; Maduro et al., 2015). Timely onset of *elt-2* appears to be critical, as its delay in early embryos has been shown to cause sustained metabolic defects in larvae despite later reattainment of wildtype ELT-2 levels (Maduro et al., 2015).

Our results suggest that END-1 mediates an all-or-none switch in the differentiation program in the absence of ELT-2 and -7. END-1 straddles both specification and differentiation, being buttressed by END-3 upstream and ELT-2/7 downstream. The only difference in these END-1 functions appears to be its timing of action and partnership with another regulatory factor. Regulatory nodes in early specification can indeed directly control morphogenetic events in various contexts (Davidson, 2010; Zhu and Rosenfeld, 2004). Interestingly, although END-1 expression decays shortly after gastrulation, we observed a persistent reduction in *act-5* expression in postembryonic larvae lacking *end-1*. One possible explanation for this observation, and the finding that patches of well-differentiated gut arise in *elt-7(-); elt-2(-)* double mutants long after END-1 becomes undetectable, might be that END-1 targets an unidentified factor (X; Figure 6) that directs differentiation in at least a subset of endodermal progenitors. Like ELT-2 and -7, expression of the putative X factor would be expected to be maintained through a positive autoregulatory loop, thereby continuing to promote gut differentiation after the expression of END-1 has subsided (Figure 6) (Sommermann et al., 2010). Alternatively, END-1 may directly activate differentiation gene batteries in early embryos, and that regulatory state might be sustained through the propagation of epigenetic memory. This priming mechanism has recently been demonstrated in the specification of the ASE sensory neurons in *C. elegans* (Charest et al., 2020). Two transiently expressed T-box factors, TBX-37 and -38, lock their target, *lsy-6*, in a transcriptionally active state during early embryogenesis, priming it for activation in restricted neuronal lineage (Charest et al., 2020). In mammals, Pax-7 (paired-box-7) initiates myogenic specification and differentiation. Interestingly, many enhancers of Pax-7 target genes retain epigenetic signatures and remain active even in the absence of the initial activator (Zhang et al., 2020).

We found that knocking down *elt-2* causes a slight but significant increase in *end-1* expression in early embryos; however, we did not observe obvious perdurance of END-1 when ELT-2 and ELT-7 are depleted, though we cannot rule out the possibility that the modest negative feedback we observed resulted from incomplete RNAi penetrance. Nevertheless, we propose that this modest feedback inhibition may function to facilitate the transition of regulatory states and ensure that developmental process moves inexorably forward. For example, repression of an early cardiac specification factor, Bmp2, by the Nkx2-5 homeodomain factor is necessary for proper morphological development of the heart in mice (Prall et al., 2007). Such transcriptional repression is also frequently used to install spatial subdivision of regulatory states (Davidson, 2010). As we have shown above, ELT-2, ELT-7 and END-1 may repress alternate cell fates in the midgut and define the boundaries of the digestive tract. It is noteworthy that structurally similar regulatory circuits are repeatedly deployed in biological networks of different contexts while performing similar functions. Thus, the functional output of a GRN depends not only on the specificity of the transcription factors, but also the underlying circuit architecture (Davidson, 2010; Peter, 2020).

### Rapid Rewiring of the Endoderm GRN in *Caenorhabditis*

How might a regulatory system of the type described here evolve? Effectors acting on terminal differentiation gene batteries, such as ELT-2 and PHA-4, are widely conserved across the animal kingdom, while the upstream inputs into GRNs appear to be recent innovations that arose during the radiation of the Elegans supergroup within the *Caenorhabditis* genus (Maduro, 2020). The *ends* and *meds* have been proposed to have arisen from the duplication of *elt-2*. Hence, gene duplication, coupled with *cis*-regulatory changes, may have led to the emergence of new circuitry and rewiring of the endoderm GRN in nematodes through intercalation of these factors in the regulatory network.

Intercalation of the MEDs and ENDs in the cascade may serve to buffer the system against environmental variation and developmental noise by freeing ELT-2 from the direct control of SKN-1, which has been shown to play conserved pleiotropic roles in stress response and lifespan regulation (reviewed in Ewe et al., 2020; Ewe et al., 2021). As we and others have shown, robust induction of *elt-2* is critical to ensure the viability and fitness of the animals (Maduro et al., 2015; Raj et al., 2010). Moreover, the deployment of MEDs and ENDs in the sequential hierarchy may allow canalization of the endodermal lineage by rapidly establishing its regulatory state in the E blastomere (Peter and Davidson, 2011). Consequently, this structure may enable increased developmental speed and early specification of the founder cells in *Caenorhabditis* species.

## Acknowledgments

We thank members of Rothman lab for helpful advice and feedback, and Cricket Wood for excellent technical support. We thank Dr. Irene Chen (University of California Los Angeles) for her helpful inputs and Dr. Zhuo Du (University of Chinese Academy of Sciences) for providing strain SYS549. We would also like to thank Dr. Rubayn Goh, Dr. Francesca Zappa, Nerea Muniozguren, and Soham Chowdhury for technical assistance. We acknowledge the use of the UCSB NRI-MCDB Microscopy Facility supported by the NIH Grant S10OD010610-01A1, as well as WormBase. Some nematode strains used in this work were provided by the Caenorhabditis Genetics Center (CGC), which is funded by NIH Office of Research Infrastructure Programs Grant P40 OD010440.

## Author contributions

Conceptualization: CKE, EMS, JHR; Resources: CKE, EMS, SEF, MFM; Formal analysis: CKE, EMS, JK; Investigation: CKE, EMS, JK; Writing - original draft: CKE; Writing – review & editing: CKE, EMS, JK, SEF, MFM, JHR; Supervision: JHR; Funding acquisition: JHR

## Data Availability

All data generated or analyzed during this study are included in this article and accompanied supplementary materials. The source code for the GRN computational model is available on https://github.com/RothmanLabCode/endoderm_GRN_model.

## Supplementary Materials

### Description of computational model

The interactions between the GATA factors (Figure 6), as well as POP-1 activation of *end-3* and *end-1*, were written as a system of differential equations. Each factor is expressed according to the concentration of its activators multiplied by respective coefficients *a*, representing the strength of activation (see Supplementary File 1). The activation is not instant, but delayed by a time interval *τ*, which accounts for the time required for transcription, translation, and re-localization to the nucleus. SKN-1 and POP-1 activation occur as square waves in EMS and E blastomere, respectively. ELT-7 and ELT-2 self-activate via feedback loops of strength *f_1_* and *f_2_*, respectively, after surpassing arbitrary concentration thresholds *Φ_1_* and *Φ_2_*, respectively. Finally, all factors are degraded at the same rate δ. The resulting system of equations is as follows.

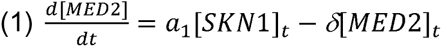

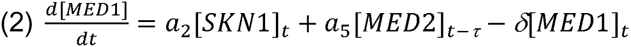

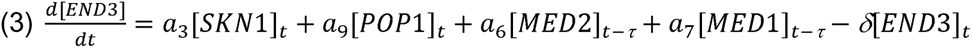

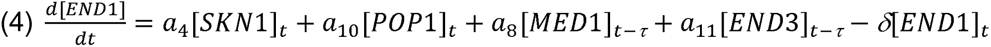

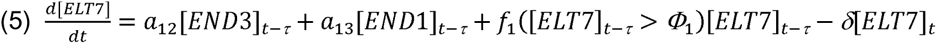

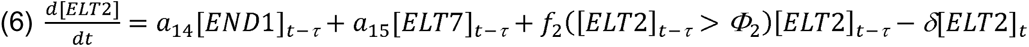

Model parameters (Supplementary File 1) were determined by fitting to transcriptomics data by a custom algorithm written in R following an iterative least-squares method. Parameter guesses were provided, and the model was run using an Euler approximation with 0.01 s time steps. The model fit was measured as the sum of squared differences between model predictions and transcriptomics data. Randomly selected parameter values were randomly changed, and the model was run again. If the resulting sum of squares was lower, the new parameter values were kept. This process was iterated ∼10^6^ times. Algorithm-selected parameter values were validated by checking model predictions against the published phenotypes of the single mutants (Boeck et al., 2011; Dineen et al., 2018; Maduro et al., 2015). Two parameters (*a_8_* and *a_14_*) were adjusted manually such that the predicted phenotype of *end-3(-)* matched that reported in published studies (Boeck et al., 2011; Maduro et al., 2005a). The source code is available on https://github.com/RothmanLabCode/endoderm_GRN_model.

**Supplementary Figure 1:**
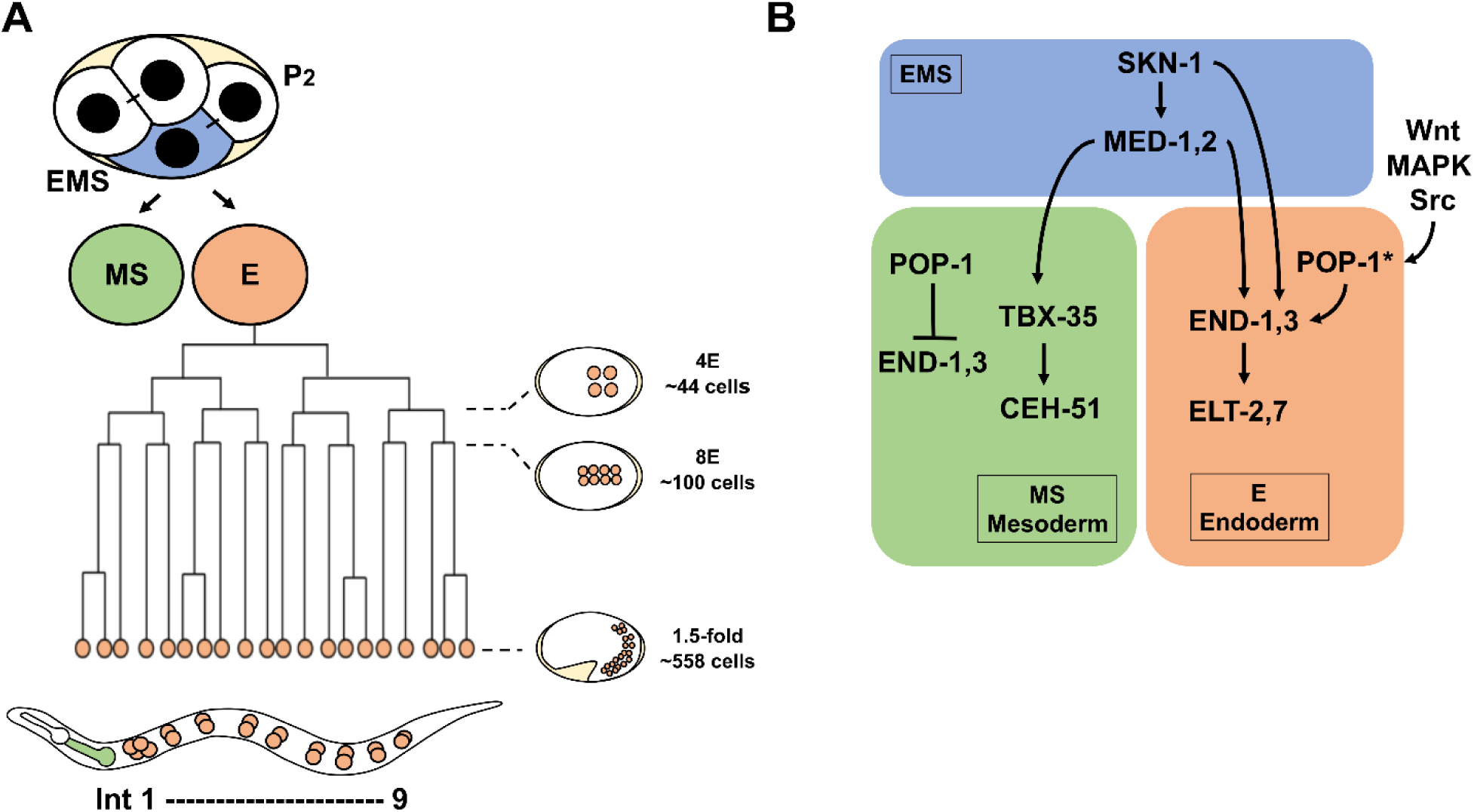
*C. elegans* mesendoderm development. (A) At the four-cell stage, EMS (blue) undergoes asymmetrical division to produce anterior MS (green) and posterior E (orange) blastomeres. Newly hatched L1s contain 20 intestinal cells arranged in nine rings (Ints). Int1 contains four cells, while the remaining eight rings contain two cells each. The MS cell gives rise mesodermal cell types, including the posterior pharynx. (B) Simplified mesendoderm GRN showing three sequential tiers of paired redundant GATA transcription factors (2 redundant MEDs 2 redundant ENDs 2 redundant ELTs). Maternally provided SKN-1 activates *med-1* and *-2*, which have both a maternal and zygotic component. In MS, POP-1 represses the expression of *end-1* and *-3* genes. MED-1 and -2 then directly activate the expression of mesoderm-specifying factor, TBX-35. In E, however, Wnt, MAPK and Src signaling from neighboring P2 cell ultimately leads to the phosphorylation of POP-1 (indicated by *), converting it from a repressor to an activator of gut fate. END-1 and -3 subsequently activate the expression of ELT-7 and -2, both of which promote gut morphological differentiation.

**Supplementary Figure 2:**
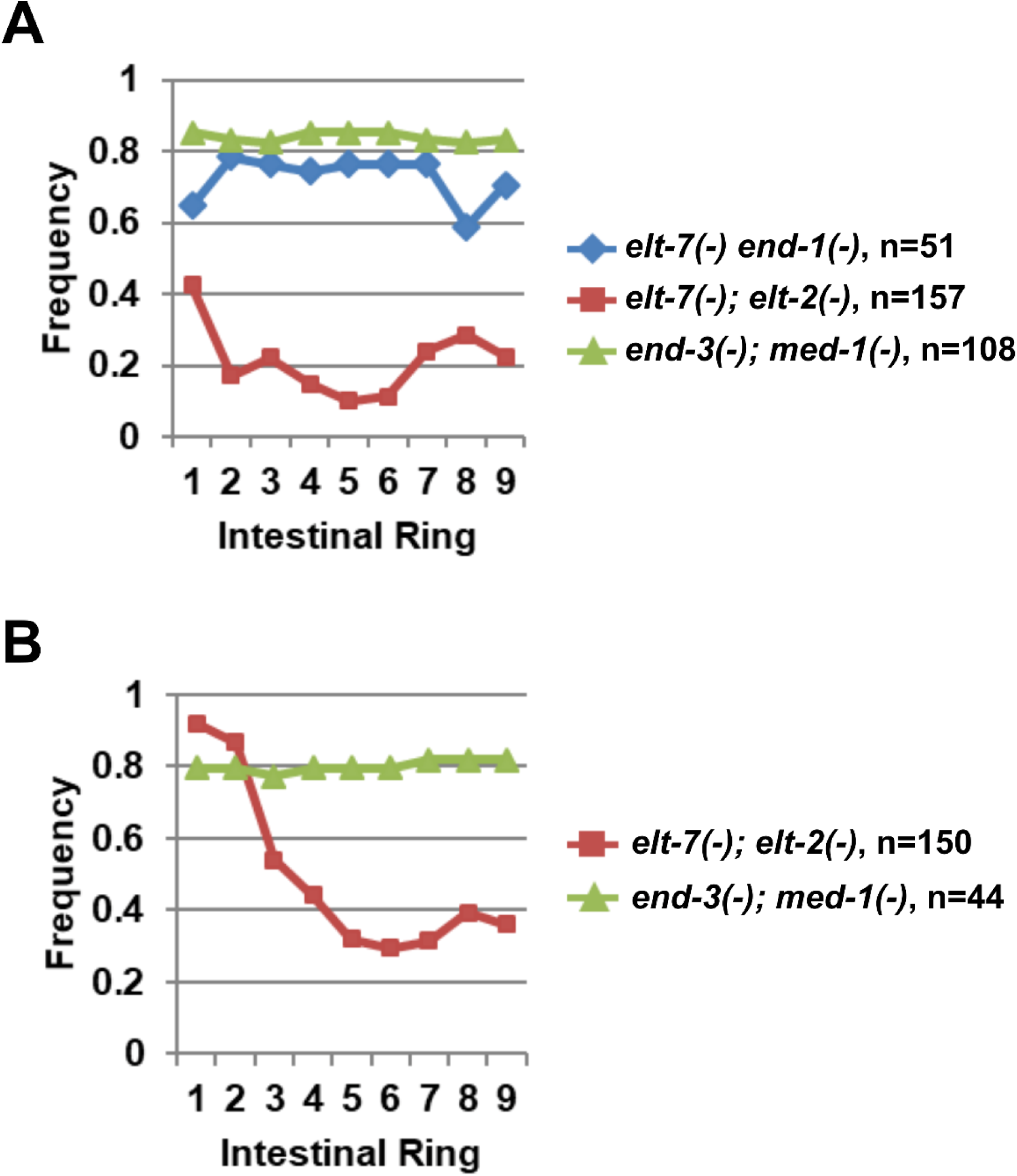
Severe gut defects in mutants lacking sequential GATA pairs. Eliminating sequential GATA pairs causes impaired gut differentiation and aberrant expression of immunoreactive (A) IFB-2 and (B) AJM-1.

**Supplementary Figure 3:**
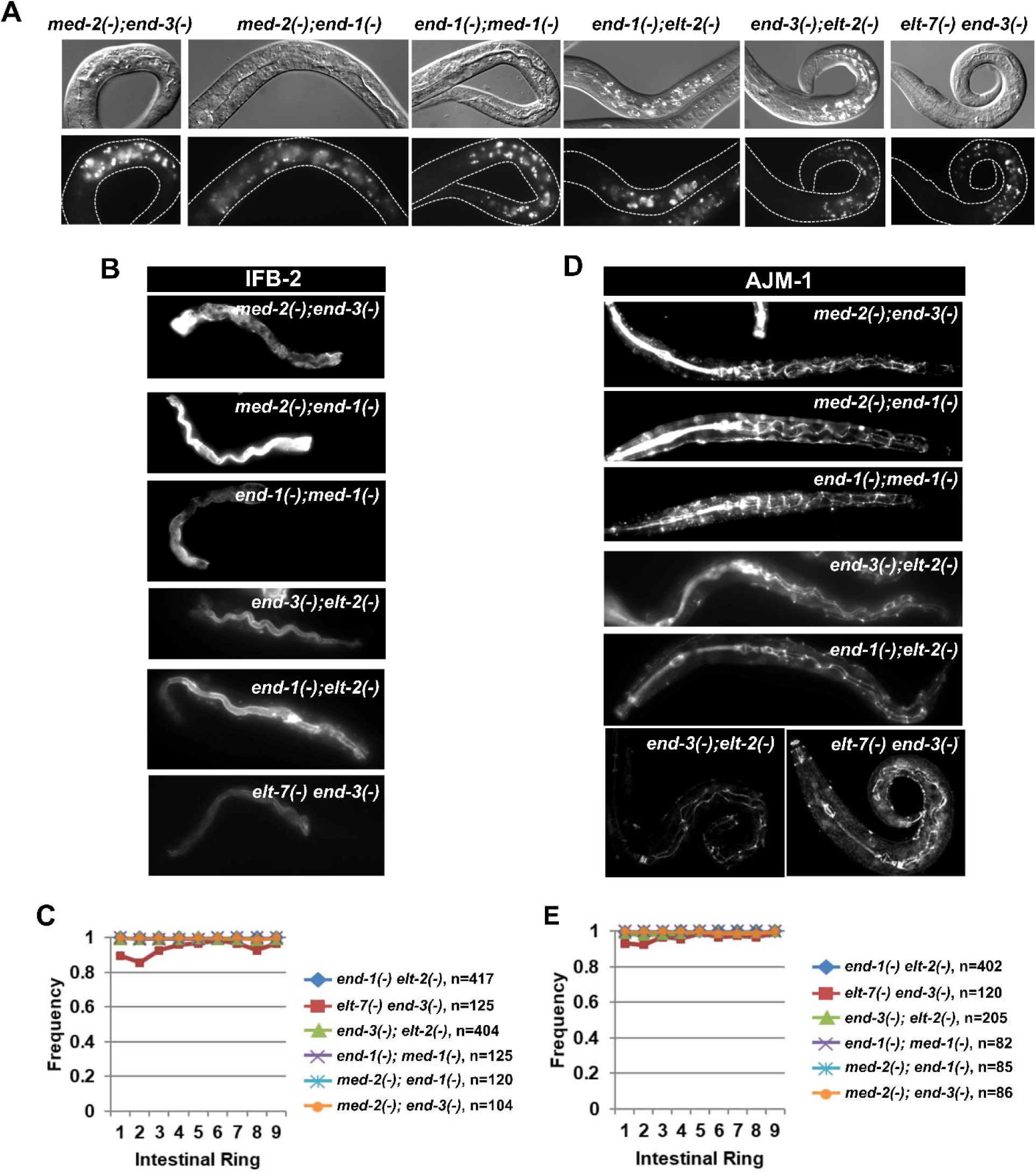
Mutants lacking alternate GATA pairs do not show apparent gut defects. (A) Mutants lacking alternate GATA pairs contain fully differentiated lumen (top row) and gut granules (bottom row) along the length of the animals. The same set of double mutants show wildtype expression of (B, C) immunoreactive IFB-2 and (D, E) AJM-1.

**Supplementary Figure 4:**
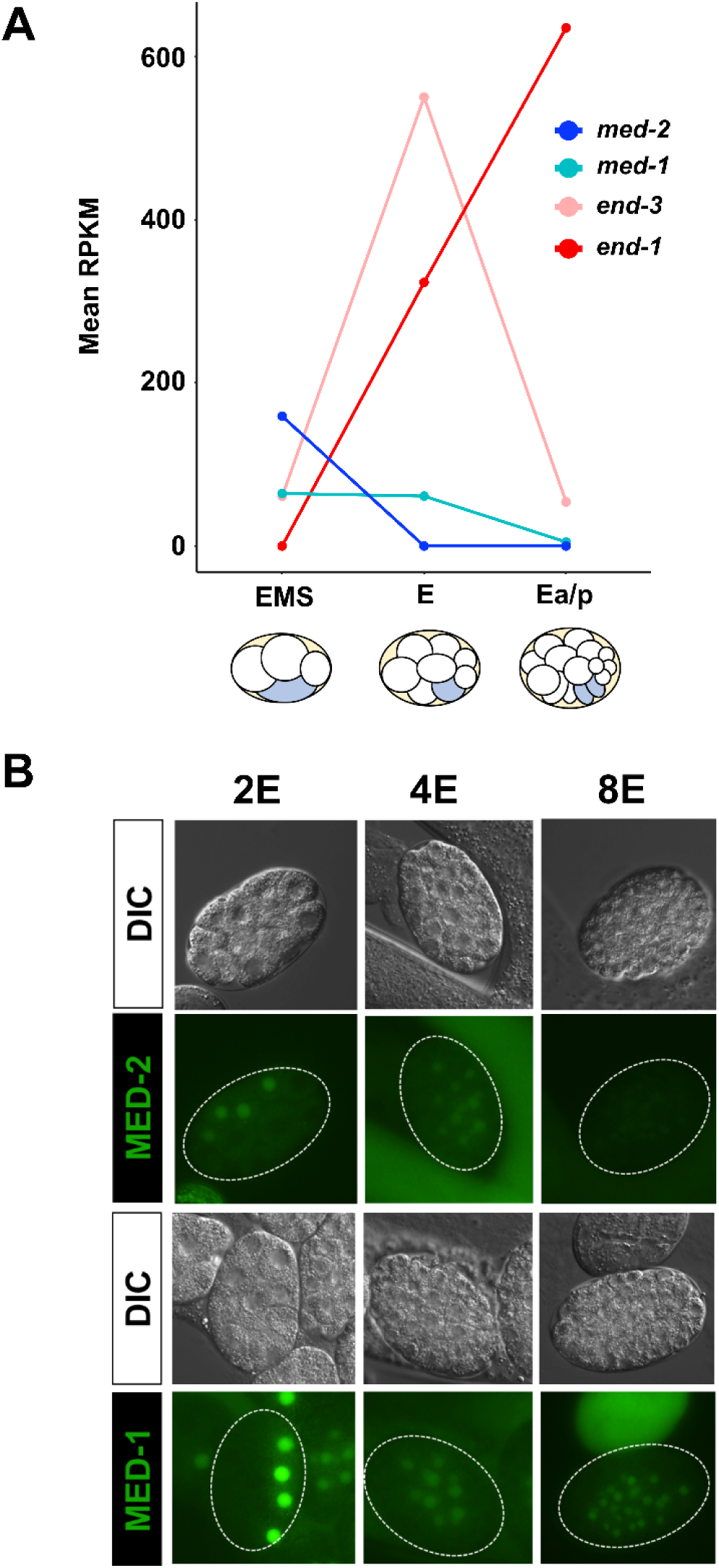
The endoderm GATA factors are deployed in temporal order. (A) Temporal expression of *med-2*, *med-1*, *end-3,* and *end-1* revealed by single-cell transcriptomic analysis (http://tintori.bio.unc.edu/) (Tintori et al., 2016). (B) Expression of MED-2 and MED-1 protein-fusion reporters in staged embryos.

**Supplementary Figure 5:**
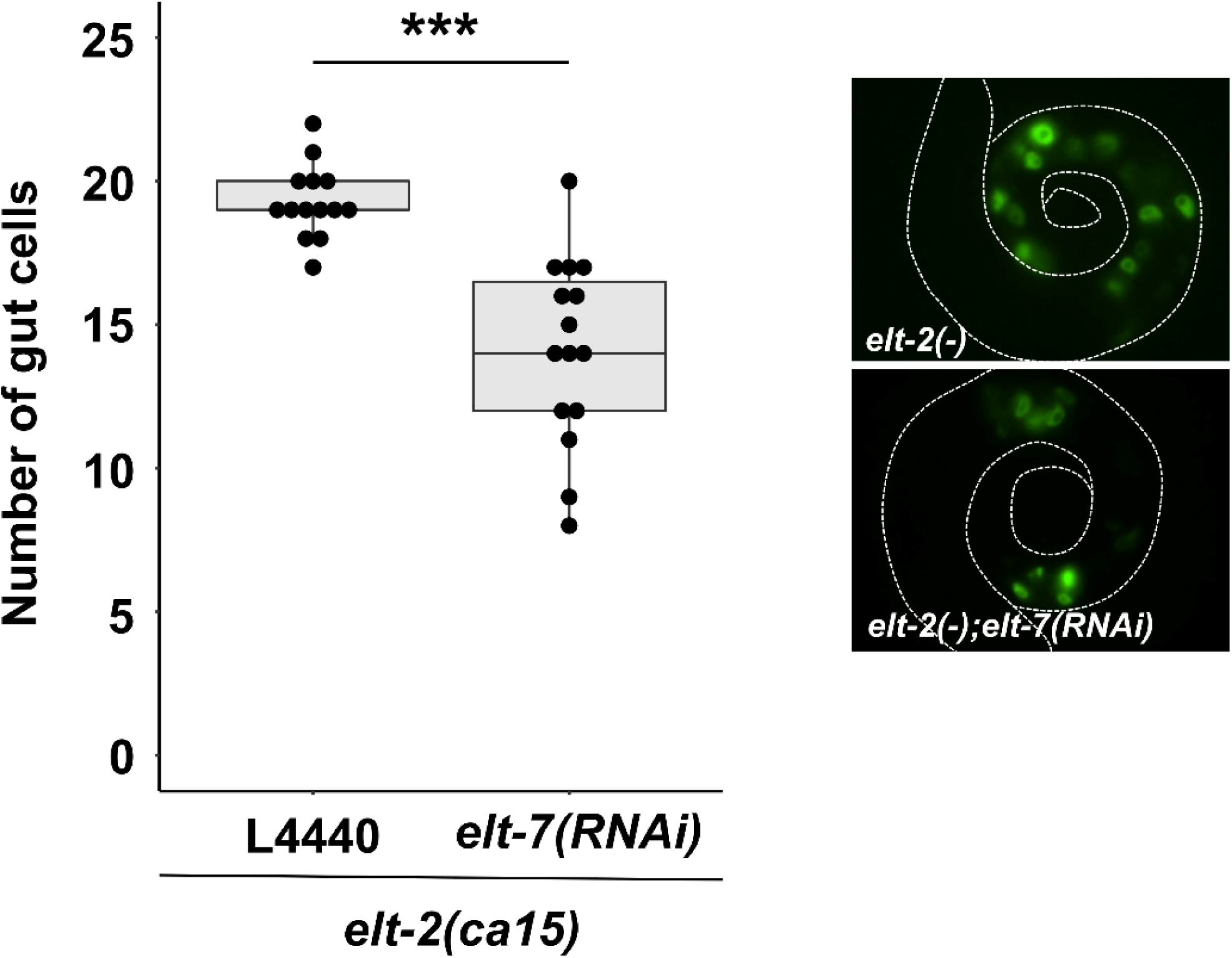
Reduced number of differentiated intestinal cells *in elt-2(-); elt-7(RNAi)* animals. On average, *elt-2(-)* animals contain 19.3 cells, while *elt-2(-); elt-7(RNAi)* animals contain 14.1 cells. The number of gut cells were scored by the expression of *elt-2p::GFP* transcriptional reporter *wIs84*. *** p ≤ 0.001 by Wilcoxon tests.

**Supplementary Figure 6:**
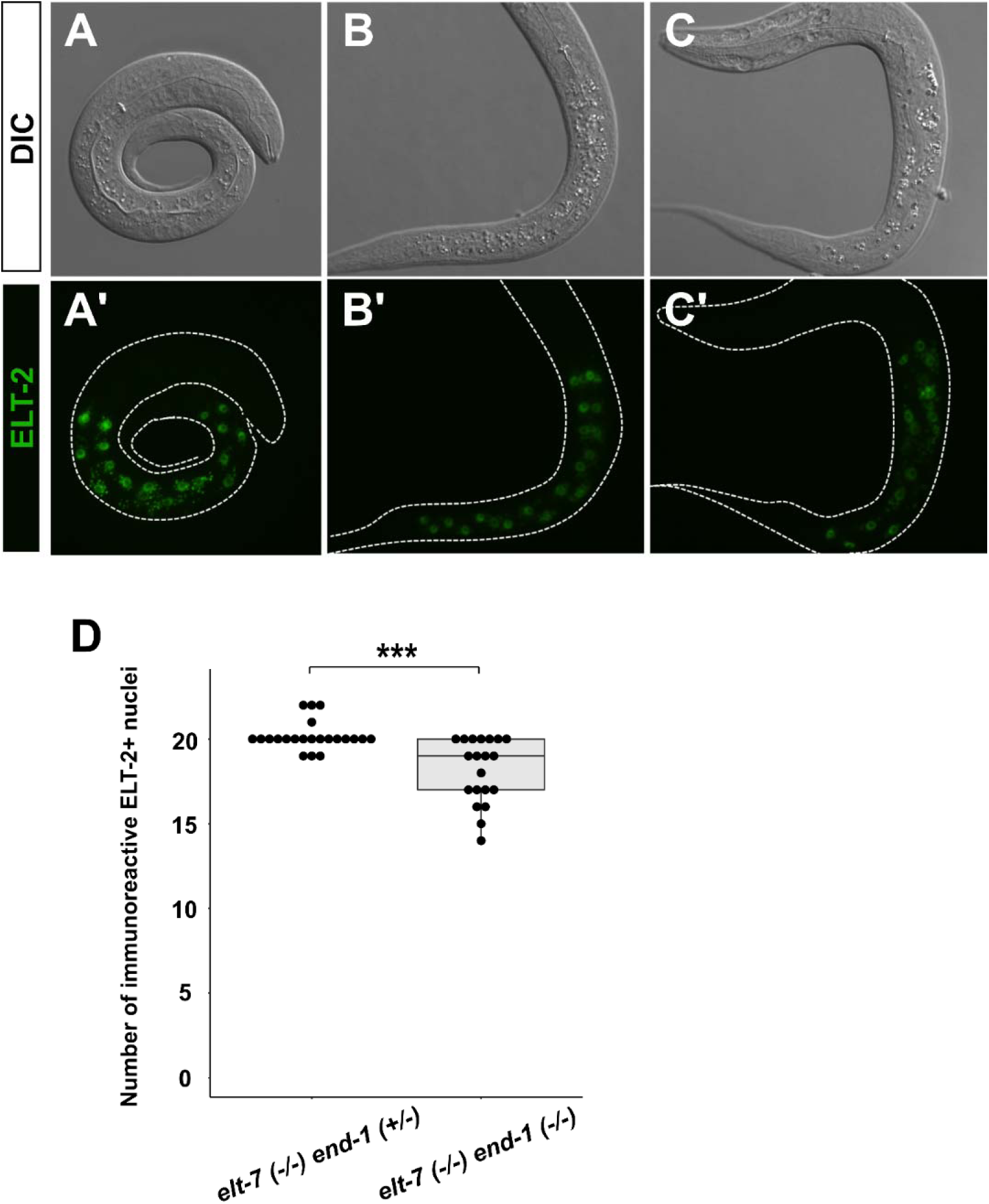
Reduced number of differentiated intestinal cells in *elt-7(-) end-1(-)* animals. (A-C) The expression of *elt-2::GFP* translational reporter in (A, A’) wildtype, (B, B’) *elt-7(-/-) end-1(+/-)*, and (C, C’) *elt-7(-/-) end-1(-/-)* L1 larvae. (D) *elt-7(-/-) end-1(-/-)* double mutants contain fewer cells expressing immunoreactive ELT-2 than *elt-7(-/-) end-1(+/-)* animals. *** p ≤ 0.001 by Wilcoxon tests.

**Supplementary Figure 7:**
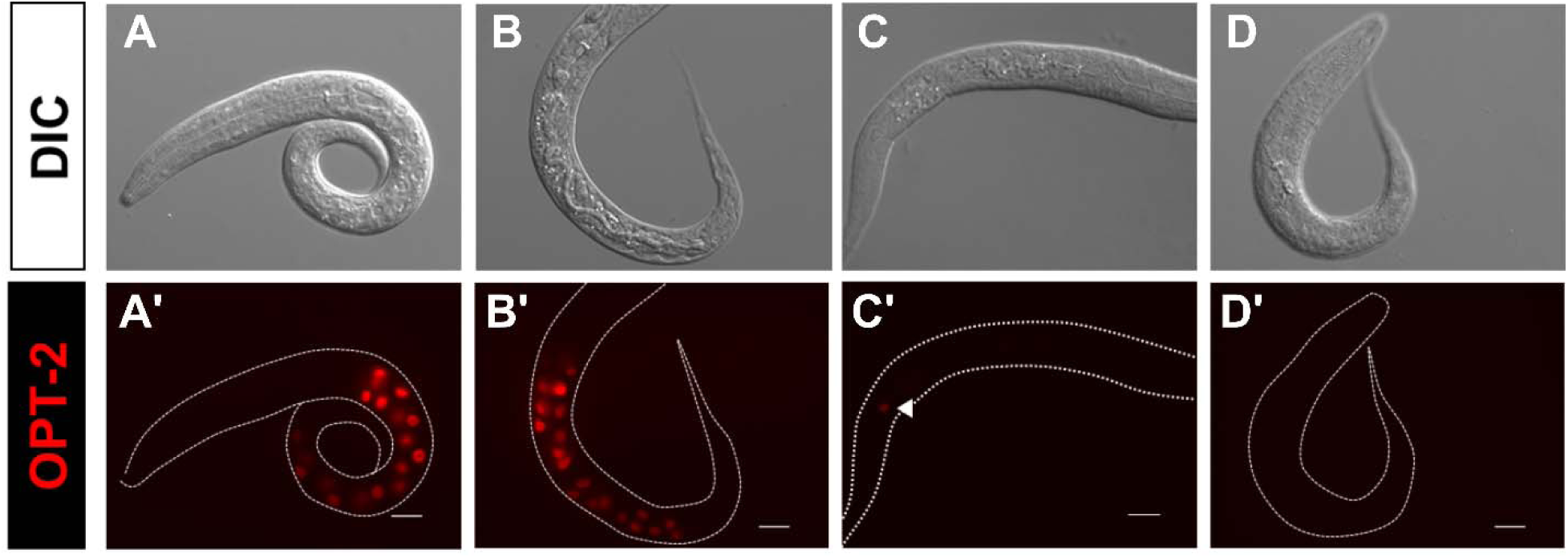
Reduced number of differentiated gut cells in differentiation-defective mutants. (A-D) DIC images of (A) wildtype, (B) *elt-2(-)*, (C) *elt-2(-); elt-7(RNAi),* and (D) *elt-7(-) end-1(-); elt-2(-)* animals. While wildtype and *elt-2(-)* worms contain a differentiated gut, *elt-2(-); elt-7(RNAi)* animals lack evident lumen and contains sporadic patches of gut granules. *elt-7(-) end-1(-); elt-2(-)* triple mutants show no apparent signs of differentiation. (A’-D’) Fluorescent images of worms in (A-D) show expression of *opt-2p::mCherry*. The number of *opt-2*-expressing cells is markedly reduced in *elt-2(-); elt-7(RNAi)* (arrowhead)*. opt-2* expression is completely abolished in *elt-7(-) end-1(-); elt-2(-).* Scale bars = 10 μm.

**Supplementary Figure 8:**
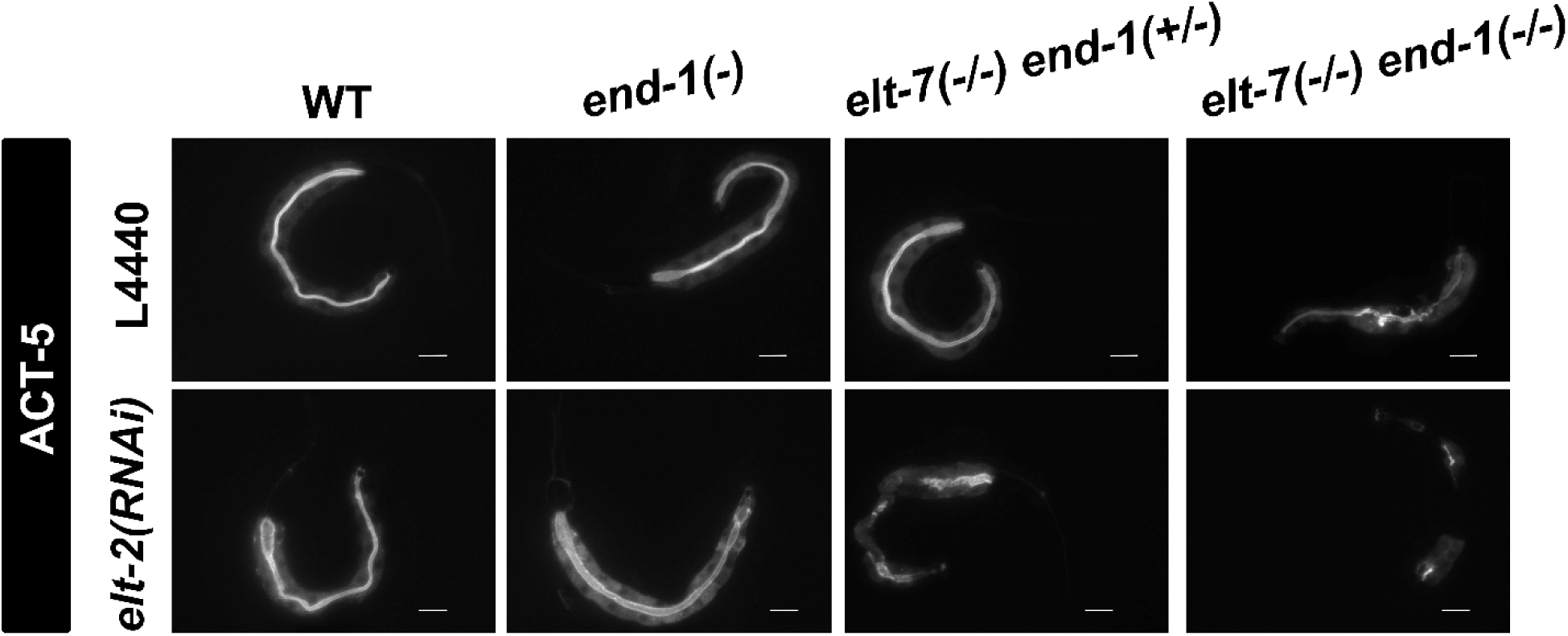
END-1, ELT-7, and ELT-2 regulate *act-5* expression. *act-5* transgene (*jyIs13*) is expressed strongly in the intestine and weakly in the excretory canal cells. *act-5::GFP* signals appear sporadic in *elt-7(-/-) end-1(+/-); elt-2(RNAi)* and *elt-7(-/-) end-1(-/-)* animals, and are almost completely missing in *elt-7(-/-) end-1(-/-); elt-2(RNAi)* mutant. The residual *act-5* expression in *elt-7(-/-) end-1(-/-); elt-2(RNAi)* may be due to incomplete RNAi penetrance. Scale bars = 10 μm.

**Supplementary Figure 9:**
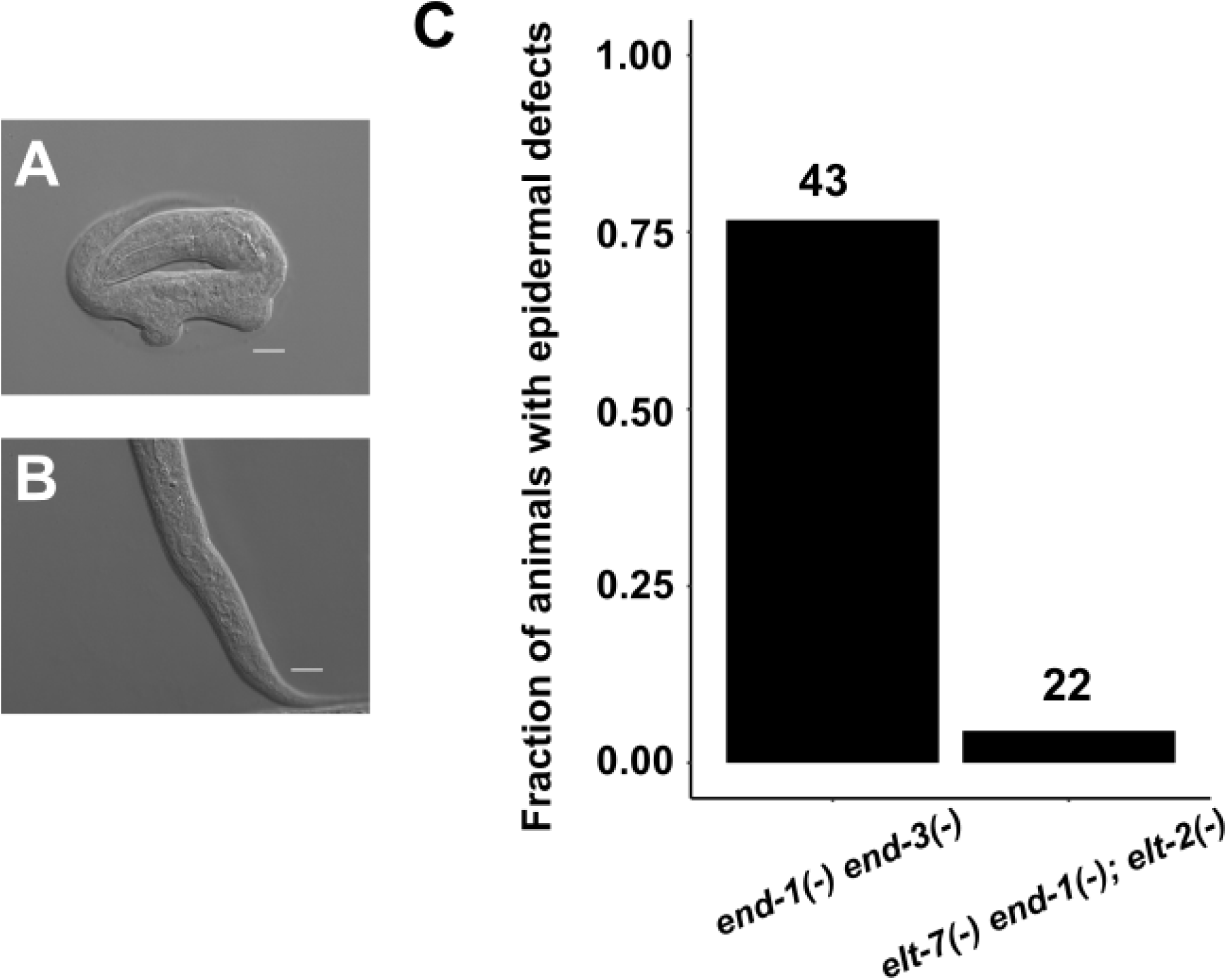
Gross epidermal defects in GATA mutants. (A) A representative *end-1(-) end-3(-)* arrested L1 shows gross epidermal defects due to E C misspecification. (B) *elt-7(-) end-1(-); elt-2(-)* mutant does not show obvious epidermal defects, as observed by DIC microscopy. Scale bars = 10 μm. (C) A large fraction of *end-1(-) end-3(-)* mutants showed deformations of the epidermis, which was rarely observed in *elt-7(-) end-1(-); elt-2(-)* worms. Number of animals scored for each genotype is indicated.

**Supplementary Figure 10:**
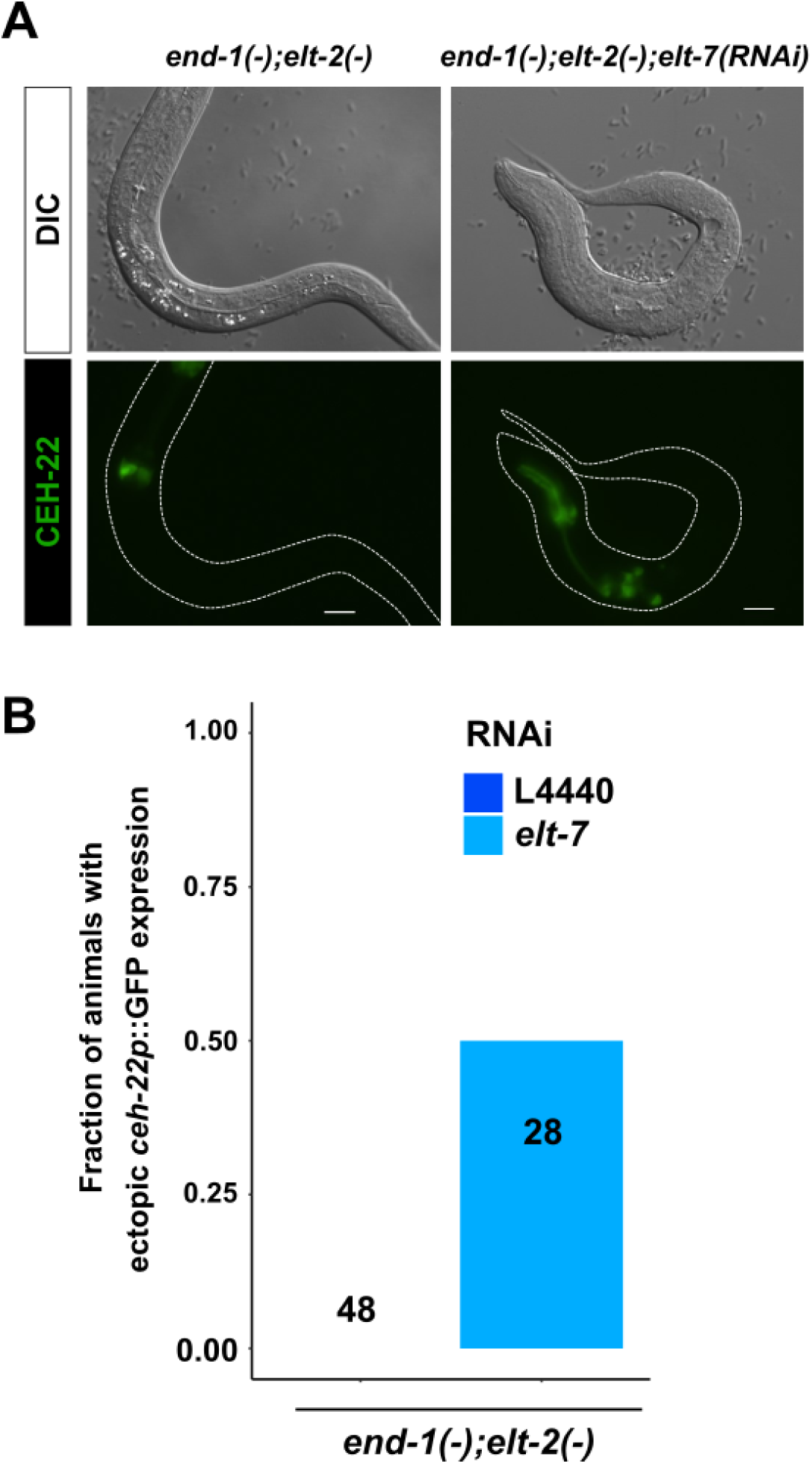
Eliminating *end-1*, *elt-7* and *elt-2* causes ectopic expression of *ceh-22* reporter. (A) A representative *end-1(-); elt-2(-)* worm contains a differentiated gut with defined lumen (top) and wildtype expression pattern of *ceh-22* that is restricted to the pharynx (bottom). On the other hand, *end-1(-); elt-2(-); elt-7(RNAi)* mutant shows no sign of gut differentiation as observed by DIC microscopy (top), and ectopic expression of *ceh-22p::GFP* reporter (bottom). Scale bars = 10 μm. (B) Knocking out *elt-7* in *end-1(-); elt-2(-)* worms causes ectopic expression of *ceh-22p::GFP* marker, as shown in (A). Number of animals scored for each genotype is indicated.

**Supplementary Table 1:** Worm strains used in this study.

| **Strain** | **Genotype** | **Source** |
| --- | --- | --- |
| MS1893 | *unc-119(ed9?) III; irSi24[opt-2p::mCherry::H2B, Cb-unc-119(+)] IV* | (Choi et al., 2017) |
| IA105 | *unc-76(e911)V; ijls12[dpy-7p::GFP::lacZ, unc-76(+)]* | (Gilleard and McGhee, 2001) |
| SYS549 | *ujls113[pie-1p::mCherry::H2B; nhr-2p::mCherry::HIS-24-let858UTR;unc119(+)] II; end-1(dev162[end-1::mNeonGreen]) V* | (Li et al., 2019) |
| JR1188 | *wls95[med-1p::GFP::MED-1+rol-6(su1006)]* | (Maduro et al., 2001) |
| MS1362 | *med-2(cx9744) III; med-1(ok804) X; irEx581[T24D3, sur-5::dsRed]* | (Maduro et al., 2007) |
| MS386 | *med-2(cx9744) III; dpy-11(e224) end-1(ok558) V* | (Maduro et al., 2007) |
| MS387 | *dpy-11(e224) end-1(ok558) V; med-1(ok804) X* | (Maduro et al., 2007) |
| MS401 | *end-3(ok1448) V; med-1(ok804) X* | (Maduro et al., 2007) |
| MS436 | *med-2(cx9744) III; end-3(ok1448) V* | (Maduro et al., 2007) |
| MS1248 | *end-1(ok558) end-3(ok1448) V; irEx568[K10F6, F58E10, pAS152(sur-5::dsRed)]* | (Owraghi et al., 2010) |
| JR3295 | *elt-7(tm840) V; elt-2(ca15) X; irEx404[unc-119::CFP and elt-2(+)]* | (Sommermann et al., 2010) |
| MS851 | *elt-2(ca15) X; irEx404[unc-119::CFP, elt-2(+)]* | (Sommermann et al., 2010) |
| RW10479 | *unc-119(ed3) III; itls37[pie-1p::mCherry::H2B::pie-1* | CGC |
|  | *3'UTR + unc-119(+)] IV; stls10116[his-72(promoter)::his-24::mCherry::let-858 3'UTR + unc-119(+)]*; *stls10426[med-2::TGF(6.2B3)::GFP::TY1::3xFLAG]* |  |
| AZ217 | *unc-119(ed3) ruls37 [myo-2p::GFP + unc-119(+)] III* | CGC |
| FX840 | *elt-7(tm840) V* | CGC |
| MS123 | *med-2(cx9744) II* | CGC |
| OP56 | *unc-119(ed3) III; gaEx290[elt-2::TY1::EGFP::3xFLAG(92C12) + unc-119(+)]* | CGC |
| RB1331 | *end-3(ok1448) V* | CGC |
| RB930 | *med-1(ok804) X* | CGC |
| VC271 | *end-1(ok558) V* | CGC |
| ERT60 | *jyls13 [act-5p::GFP::ACT-5 + rol-6(su1006)] II* | CGC |
| OH15876 | *pha-4(ot946[pha-4::GFP]) V* | CGC |
| JR4405 | *ruls37 [myo-2p::GFP + unc-119(+)] III; him-5(e1490) V; elt-2(ca15) X; IrEx404[unc-119::CFP, elt-2(+)]* | This study |
| JR4384 | *ruls37 [myo-2p::GFP + unc-119(+)] III; elt-7(tm840) V; elt-2(ca15) X; IrEx404[unc-119::CFP, elt-2(+)]* | This study |
| JR4369 | *hum-1(cv21[hum-1::RFP]) I; elt-7(tm840) end-1(ox134)/ elt-7(tm840) tmC12[egl-9(tmls1197)] V; gaEx290[elt-2::TY1::EGFP::3xFLAG(92C12) + unc-119(+)]* | This study |
| JR4371 | *hum-1(cv21[hum-1::RFP]) I; gaEx290[elt-2::TY1::EGFP::3xFLAG(92C12) + unc-119(+)]* | This study |
| JR4368 | *ruls37 [myo-2p::GFP + unc-119(+)] III; end-1(ok558) end-3(ok1448) V; irEx568[K10F6, F58E10, pAS152(sur-5::dsRed)]* | This study |
| JR2488 | *him-8(e1489) IV; end-1(ox134) V* | This study |
| JR3341 | *elt-7(tm840) end-1(ox134)/ dpy-11(e224) unc-76(e911) V* | This study |
| JR3342 | *elt-7(tm840) end-3(ok1448) V* | This study |
| JR4037 | *elt-2(ca15) X; iJls12[dpy-7p::GFP::lacZ, unc-76(+)]; irEx404[unc-119::CFP, elt-2(+)]* | This study |
| JR4063 | *him-8(e1489) irSi24[opt-2p::mCherry::H2B, Cb-unc-119(+)] IV; elt-2(ca15) X; irEx404[unc-119::CFP, elt-2(+)]* | This study |
| JR4097 | *elt-7(tm840) end-1(ox134)/ elt-7(tm840) tmC12[egl-9(tmls1197)] V* | This study |
| JR4194 | *jcls1[ajm-1p::AJM-1::GFP + unc-29(+)+ rol-6(su1006)] IV; elt-7(tm840) end-1(ox134)/ elt-7(tm840) tmC12[egl-9(tmls1197)] V* | This study |
| JR4195 | *elt-7 (tm840) end-1(ox134)/ elt-7(tm840) tmC12[egl-9(tmls1197)] V; elt-2(ca15) X; irEx404[unc-119::CFP, elt-2(+)]* | This study |
| JR4200 | *him-8(e1489) irSi24 [opt-2p::mCherry::H2B, Cb-unc-119(+)] IV; elt-7(tm840) end-1(ox134)/ elt-7(tm840) tmC12[egl-9(tmls1197)] V; elt-2(ca15) X; irEx404[unc-119::CFP, elt-2(+)]* | This study |
| JR4304 | *elt-7 (tm840) end-1(ox134)/ elt-7(tm840) tmC12[egl-9(tmls1197)] V; wEx1751[cdf-1p::CDF-1::GFP + rol-6(su1006)]* | This study |
| JR4313 | *elt-7(tm840) end-1(ox134)/ elt-7(tm840) tmC12[egl-9(tmls1197)] V; gaEx290[elt-2::TY1::EGFP::3xFLAG(92C12) + unc-119(+)]* | This study |
| JR4364 | *ruls37 [myo-2p::GFP + unc-119(+)] III; elt-7(tm840) end-1(ox134)/ elt-7(tm840) tmC12[egl-9(tmls1197)] V; elt-2(ca15) X; irEx404[unc-119::CFP, elt-2(+)]* | This study |
| JR4367 | *ruls37 [myo-2p::GFP + unc-119(+)] III; elt-7(tm840) end-1(ox134)/ elt-7(tm840) tmC12[egl-9(tmls1197)] V* | This study |
| MS880 | *dpy-11(e224) end-1(ok558) V; elt-2(ca15) X;* | This study |
|  | *irEx404[unc-119::CFP, elt-2(+)]* |  |
| JR4407 | *jcls1[ajm-1p::AJM-1::GFP + unc-29(+)] + rol-6(su1006)] IV; end-1(ox134) V* | This study |
| MS900 | *end-3(ok1448) V; elt-2(ca15) X; irEx404[unc-119::CFP, elt-2(+)]* | This study |
| JR4412 | *jyls13 [act-5p::GFP::ACT-5 + rol-6(su1006)] II; elt-7(tm840) end-1(ox134)/ elt-7(tm840) tmC12[egl-9(tmls1197)] V* | This study |
| JR4413 | *jyls13 [act-5p::GFP::ACT-5 + rol-6(su1006)] II; end-1(ox134) V* | This study |
| JR4439 | *him-8(e1489) IV; culs1[ceh-22p::GFP + rol-6(su1006)] end-1(ox134) V; elt-2(ca15); irEx404[unc-119::CFP, elt-2(+)]* | This study |
| JR4446 | *elt-7(tm840) end-1(ox134)/ elt-7(tm840) tmC12[egl-9(tmls1197)] V; kcls6[ifb-2p::IFB-2::CFP]* | This study |
| JR4198 | *end-1(ok558) end-3(ok1448) V; ijls12[dpy-7p::GFP::lacZ, unc-76(+)]; irEx568[K10F6, F58E10, pAS152(sur-5::dsRed)]* | This study |
| JR4452 | *end-1(ox134) V; elt-2(ca15); ijls12[dpy-7p::GFP::lacZ, unc-76(+)]; irEx404[unc-119::CFP, elt-2(+)]* | This study |
| JR3315 | *elt-2(ca15) X; wls84[elt-2p::GFP+ rol-6(su1006)]; irEx404[unc-119::CFP, elt-2(+)]* | This study |
| JR4463 | *pha-4(ot946[pha-4::GFP]) V; elt-2(ca15) X; irEx404[unc-119::CFP, elt-2(+)]* | This study |

**Supplementary File 1 (.xlsx)**: Model parameters and outputs of the endoderm GRN. Expression of each factor is determined by the concentration of its activators multiplied by a coefficient *a*, representing the strength of the inputs. SKN-1 expression follows a square wave in EMS blastomere. POP-1 expression follows a square wave in the E cell. Feedback coefficients *f* become nonzero once their respective factors surpass a certain threshold *Φ*.

